# A self-blinking DNA probe for 3D superresolution imaging of native chromatin

**DOI:** 10.1101/2022.06.22.497271

**Authors:** Yang Zheng, Sen Ye, Shumin Li, Cuifang Liu, Shihang Luo, Ruiqin Xia, Yanqin Chen, Yunsheng Li, Zhenni Zhu, Lingyi Huang, Shan Deng, Karen Wing Yee Yuen, Ping Chen, Yongdeng Zhang, Wei Ji, Ruibang Luo, Guohong Li, Dan Yang

## Abstract

Single-molecule localization microscopy is a powerful superresolution imaging technique to study biological questions by visualizing subcellular fine structures with nanometer-scale precision. However, its application in live-cell imaging studies has been impeded by the paucity of self-blinking organic fluorophores that enable high spatiotemporal resolution and labeling/localization density at a moderate laser intensity. Herein, we report a self-blinking Si-rhodamine dye **6-HESiR** with a suitably increased “ON” fraction and a fluorogenic self-blinking dsDNA probe **6-HoeHESiR** as a powerful tool for 3D superresolution imaging of native chromatin in eukaryotes without the use of photoswitching buffer and high laser intensity. With the probe **6-HoeHESiR**, 3D superresolution imaging of in vitro reconstituted nucleosomal arrays and chromatin fibers yielded results consistent with EM analysis. Similar euchromatin and heterochromatin structures were visualized in fixed and live cells with high spatiotemporal resolution and labeling density, providing the first live-cell evidence for a hierarchical model of chromatin organization. 3D imaging results obtained in the presence of selective inhibitors of histone deacetylases also corroborate chromatin fiber decompaction upon hyperacetylation of histones.

## INTRODUCTION

Superresolution fluorescence microscopy is a powerful tool for interrogating important biological questions. By bypassing the diffraction limit of light, superresolution microscopy has been developed to visualize previously unresolvable fine subcellular structures with nanometer-scale precision^1–6^. However, the best-performing commercial organic dyes (e.g., Alexa 647) widely used for single-molecule localization microscopy (SMLM) still suffer from poor cell permeability and the need for photoswitching buffers that are highly reducing and incompatible with live cells^2,6^. Such limitations have impeded the study of important biological questions by live-cell SMLM. In recent years, self-blinking organic fluorophores such as **HMSiR**^7–10^ were developed for SMLM of living cells without using live-cell incompatible photoswitching buffers and 405-nm laser. Yet, high-quality imaging results with high spatiotemporal resolution and labeling density were only achieved in 2D imaging of endoplasmic reticulum (ER)^11^ with very high laser intensity^12^. It is still a formidable challenge to achieve high spatiotemporal resolution and labeling density in 3D live-cell SMLM, even at a moderate excitation laser intensity. In principle, the number of “ON” state fluorophores in each frame should be as high as possible to minimize the acquisition time, yet low enough to avoid PSF overlaps^6^. In this work, we have developed a self-blinking Si-rhodamine dye **6-HESiR** with a suitably increased “ON” fraction by employing the structural fine-tuning strategy^13–15^. By connecting **6-HESiR** to a Hoechst fragment for DNA minor groove binding, we have obtained a self-blinking fluorogenic probe **6-HoeHESiR** for imaging native chromatin in living cells. This probe has been validated with lambda DNA, *in vitro* reconstituted nucleosomal arrays and 30-nm chromatin fibers, and our 3D imaging results are consistent with those obtained by EM analysis. The 3D visualization of both nucleosomal arrays and chromatin fibers in fixed cells and living cells has been achieved with high spatiotemporal resolution and labeling density at low laser intensity to minimize phototoxicity. We have observed structural plasticity of chromatin fibers in both HeLa cells and chicken erythrocytes. Fast chromatin fiber dynamics were captured with less than 2-second temporal resolution, and the mean resting/dwelling lifetime of chromatin fibers was estimated to be 0.6 seconds in HeLa cells and 1 second in chicken erythrocytes. We have also observed the decompaction of chromatin fibers induced by small-molecule inhibitors of histone deacetylases (HDACs), providing the first live-cell evidence for the direct link between histone acetylation state and 3D chromatin organization with unprecedented resolution.

## RESULTS

### The development of a self-blinking fluorogenic DNA probe

To achieve high spatiotemporal resolution and labeling density in 3D live-cell SMLM, we conceived a design strategy for self-blinking fluorophores with the following criteria: 1) absorption maximum in the deep red and near infrared regions to minimize autofluorescence and phototoxicity to living cells^12^; 2) a fine-tuned fluorescent “ON” state population for optimal localization number per frame to achieve sufficient labeling density in a short acquisition time while preventing PSF overlap^6^; 3) take into account the pH and hydrophobicity that influence the “ON” state population^11^. To meet the above criteria, we developed a self-blinking Si-rhodamine dye **6-HESiR** (Fig. 1a-b) by replacing the hydroxymethyl group of **HMSiR^7^** with a hydroxyethyl group^16,17^. This type of self-blinking dyes is expected to undergo equilibrium between the open form (fluorescent “ON” state) and the closed form (“OFF” state) with an equilibrium constant as *K*cycl. For entropic reason, formation of the six-membered-ring cyclic form (‘OFF” form) for **6-HESiR** should be more difficult than that of the five-membered-ring cyclic form of **HMSiR**. As a result, **6-HESiR** should favor the open form more than **HMSiR**, and hence increasing the localization number per frame significantly for fast acquisition of sufficient localizations to resolve dynamic fine structures. This new fluorophore **6-HESiR** was successfully prepared through an efficient synthetic route (shown in Supplementary Information), and the “ON” state population of **6-HESiR** was estimated from the absorption spectra to be 20% at physiological pH (7.4) in aqueous buffer with **SiR650** as the benchmark of the 100% “ON” state (Fig. 1b-c, Supplementary Fig. 1). In addition, based on the pH titration curve (Fig. 1d), we determined the p*K*cycl value of **6-HESiR** to be 6.6, higher than that of **HMSiR** (p*K*cycl = 5.8, Fig. 1a).

**Fig. 1.**
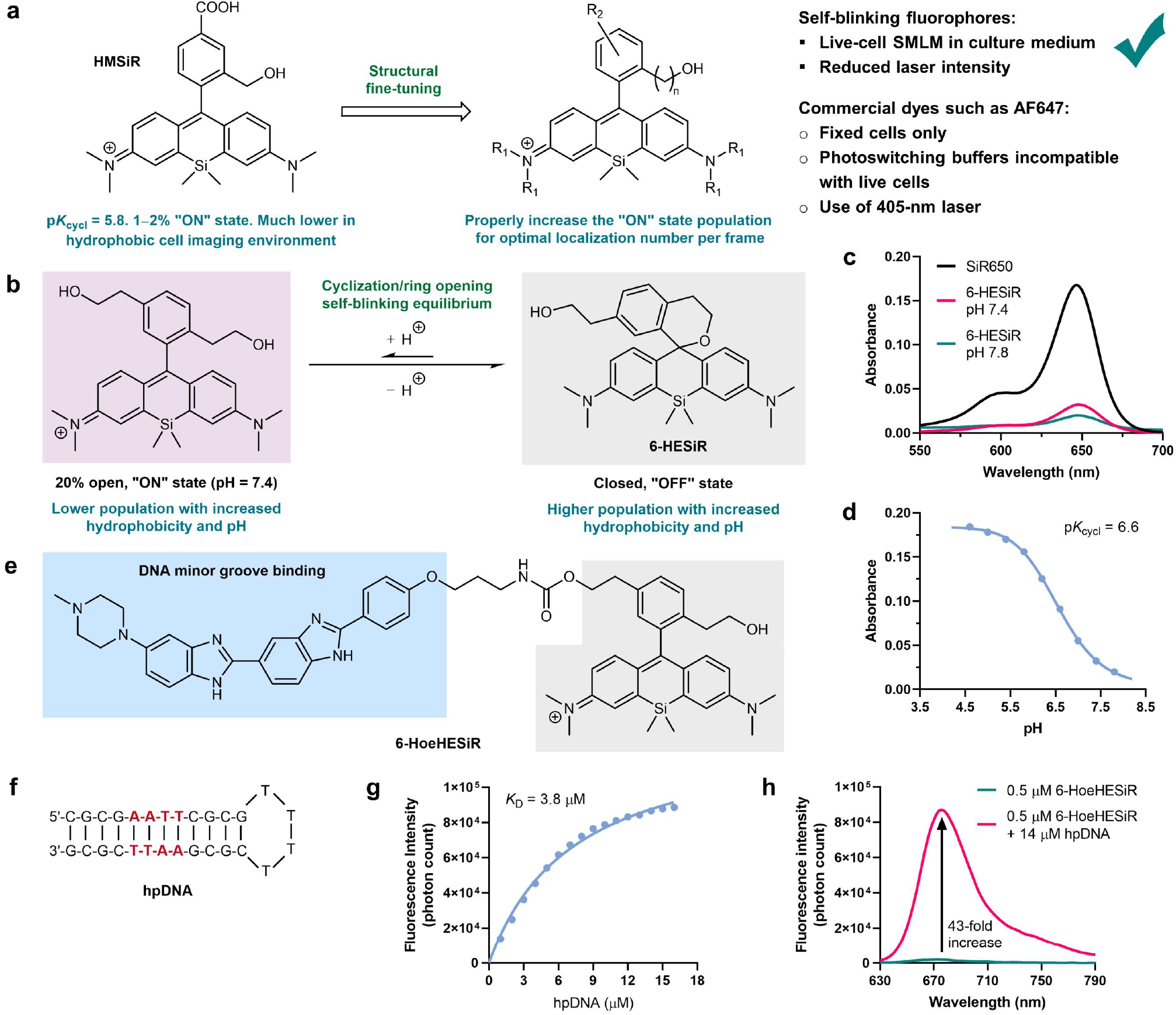
Development of 6-HESiR and 6-HoeHESiR. **a**, Structural fine-tuning strategy for developing self-blinking fluorophores. **b**, Self-blinking mechanism. **c**, Absorption spectrum of **6-HESiR** (2 μM) in 0.1 M potassium phosphate buffer (pH = 7.4 and 7.8) with **SiR650** (2 μM) as the “100%” benchmark. Abs_max_ at 648 nm. Fl_max_ at 665 nm. **d**, pH titration of **6-HESiR** (2 μM) in 0.1 M potassium phosphate buffer. **e**, Design of **6-HoeHESiR** by Hoechst tagging strategy. **f**, The hpDNA for fluorescence titration. **g**, Fluorescence titration of **6-HoeHESiR** (0.5 μM) with hpDNA (0–16 μM) in Tris-HCl saline buffer (50 mM Tris-HCl, 100 mM NaCl, pH = 7.4). Fl_max_ at 676 nm. **h**, Fluorescence increases 43-fold in the presence of 14 μM hpDNA.

The successful implementation of live-cell SMLM hinges not only on the self-blinking fluorophore but also on a fully biocompatible labeling strategy that is noninvasive, highly selective, and artifact-free with high labeling density. In this regard, we chose the Hoechst tagging strategy^18^, as it has found some success in developing fluorogenic probes for live-cell chromatin imaging^18–23^ with low cytotoxicity. The Hoechst moiety could noncovalently bind to the DNA minor groove in a highly selective manner. Therefore, we designed and synthesized **6-HoeHESiR** by connecting **6-HESiR** and the Hoechst fragment with a carbamate linker (Fig. 1e). The *in vitro* binding of **6-HoeHESiR** to a benchmark hairpin DNA (hpDNA) molecule (Fig. 1f) with an AATT Hoechst binding site^24^ was examined by fluorescence titration. In the absence of hpDNA, the fluorescence of **6-HoeHESiR** was nearly quenched (Fig. 1h). A dose-dependent fluorescence turn-on was observed upon the addition of increasing equivalents of hpDNA to **6-HoeHESiR** (Fig. 1g). A 43-fold fluorescence increase (Fig. 1h) with 14 μM of hpDNA confirmed the sensitivity and fluorogenicity of **6-HoeHESiR**. Based on the titration curve (Fig. 1g), the dissociation constant (*K*_D_) of the complex formed by **6-HoeHESiR** and hpDNA was estimated to be 3.8 μM. The attachment of **6-HESiR** to the Hoechst dye caused a slight redshift of both the absorbance and fluorescence maxima of **6-HoeHESiR** (Supplementary Fig. 2a-b). Upon the addition of hpDNA (14 μM), the absorbance of **6-HoeHESiR** was increased to a level close to that of the free fluorophore **6-HESiR** (Supplementary Fig. 2a), and the fluorescence of **6-HoeHESiR** was recovered to 21% of that of **6-HESiR** (Supplementary Fig. 2b).

### Validation with lambda DNA and *in vitro* reconstituted chromatin structures

To evaluate our new probe **6-HoeHESiR**, we first attempted an SMLM experiment of spin-coated lambda DNA^25^ with (A/T)_4_ binding sites for the Hoechst moiety^24^. A full width at half maximum (FWHM) of 26 nm was successfully obtained (Supplementary Fig. 3), which is within the range of 20–30 nm as reported in literature^25^. Then, we focused on *in vitro* reconstituted nucleosomal arrays and 30-nm chromatin fibers as benchmarks previously well characterized by EM^26^ (Fig. 2d and 2f, Supplementary Fig. 4a and 4e). Apart from the AATT binding site, three additional Hoechst binding sites^24^ in the DNA template for *in vitro* reconstituted chromatin samples were also identified by fluorescence titration (Fig. 2a, Supplementary Fig. 2c-f). Based on the dissociation constants, the half-life of the complex formed by **6-HoeHESiR** and a nucleosome was estimated to be 10–20 ms^24^. Thus, fast binding/dissociation cycles between **6-HoeHESiR** and nucleosomes could allow fast self-blinking with a pool of **6-HoeHESiR** molecules in the unbound state (Fig. 2b). Since the hydrophobic environment in the nucleus favors the closed “OFF” state^11^ (Fig. 1b) and the open form of **6-HoeHESiR** in the unbound state is nearly quenched in fluorescence (Fig. 1h), the “ON” state population of **6-HoeHESiR** in chromatin imaging is expected to be lower than the 20% “ON” fraction of **6-HESiR** measured in aqueous buffer. Gratifyingly, **6-HoeHESiR** exhibited excellent self-blinking in SMLM experiments on reconstituted chromatin samples at a modest laser intensity^12^ of 950 W/cm^2^ (Supplementary Video 1). The reconstituted nucleosomal arrays displayed the pattern of discrete fluorescence signals (Fig. 2c and 2e, Supplementary Fig. 4b), which resembles the EM images (Fig. 2d, Supplementary Fig. 4a). Analysis of localizations on individual nucleosomes indicated that the ratio of axial to lateral resolution is 2.3 (Supplementary Fig. 4c-d). To extract 3D structural features of reconstituted 30-nm chromatin fibers visualized in 3D SMLM experiments (Fig. 2c and 2g; Supplementary Fig. 4f), we developed an image analysis program (Supplementary Fig.5 and 6, Supplementary Information). Statistical results showed that the reconstituted chromatin fibers had a mean diameter of 32 nm, length of 108 nm, mean localization precision of 7.8 nm and labeling density of 31,815 molecules/μm^3^ (Fig. 2h, Supplementary Fig. 4g, Supplementary Video 2 and 3). Although the reconstituted 30-nm fibers without biotin labels were visualized by EM in a dry and flattened state (Supplementary Information), unlike our 3D SMLM approach, the 3D structural features obtained with **6-HoeHESiR** were consistent with those revealed by EM (Fig. 2f-h). Some elongated fibers were observed, probably due to slight loosening in the aqueous buffer without chemical crosslinker compared with the dry and crosslinked state in EM analysis.

**Fig. 2.**
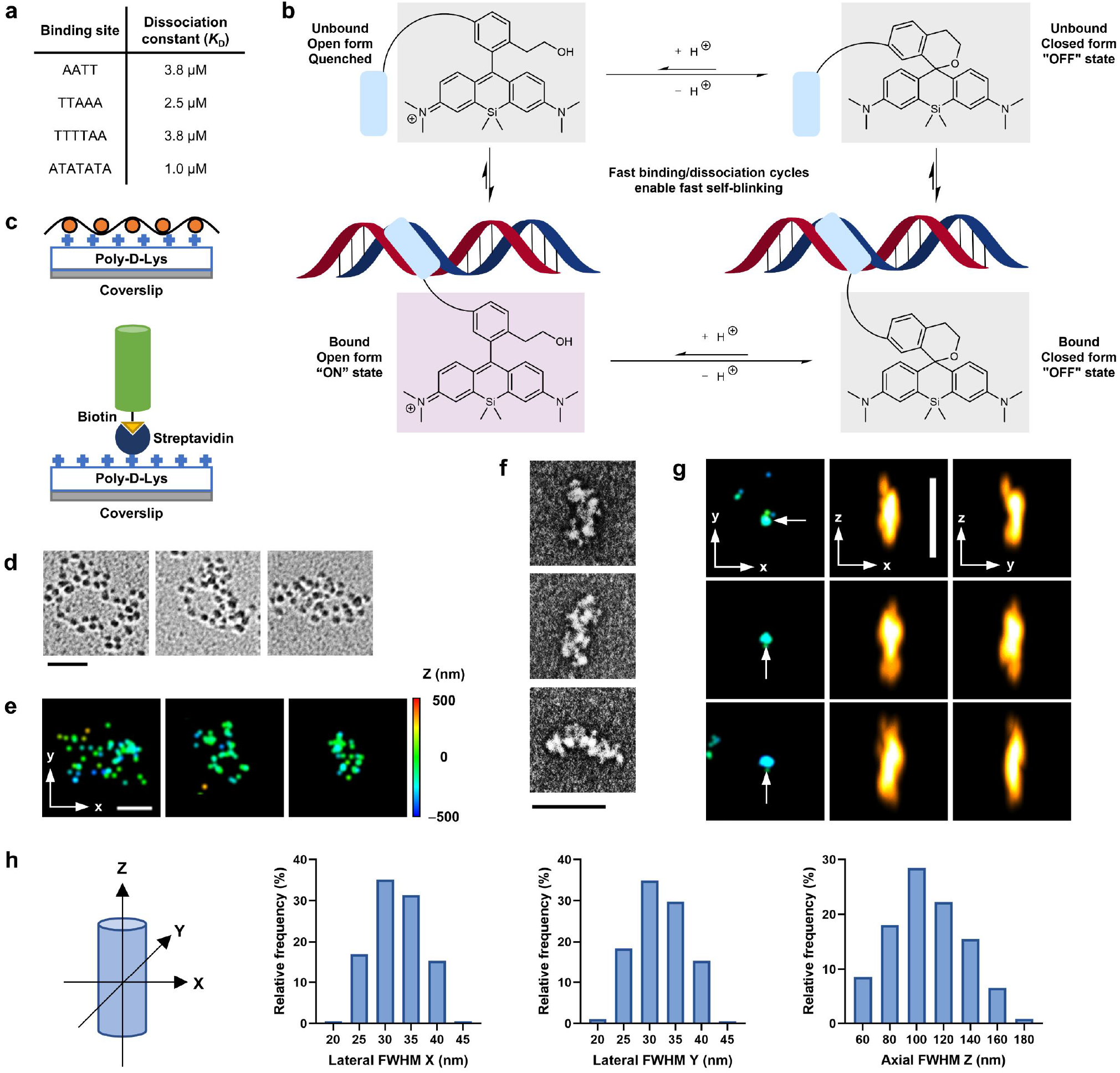
in vitro reconstituted chromatin structures. **a**, Dissociation constants of four binding sites in the DNA template. **b**, Working mechanism of **6-HoeHESiR** in the presence of nucleosomes. **c**, Immobilization approaches for 40 × 187-bp nucleosomal arrays and 30-nm chromatin fibers. **d**, EM images of nucleosomal arrays. **e**, Images of nucleosomal arrays by 3D SMLM. **f**, EM images of 30-nm chromatin fibers. **g**, Images of 30-nm chromatin fibers by 3D SMLM. 3D sizes by FWHM (x, y, z) in nm (from top to bottom): 29, 30, 93; 30, 30, 101; 34, 26, 104. Localization precision (nm): 5.5, 5.9, 6. **h**, Histograms of 3D sizes of chromatin fibers. *N* = 370 from 3 fields of view. Incubation with **6-HoeHESiR** (2.5 μM) for 10 min. Scale bar: 100 nm in **d** and **f**; 200 nm in **e** and **g**. Laser intensity: 0.95 kW/cm^2^ at 656 nm. Axial position (nm) represented by RGB color depth coding.

### Native chromatin 3D structures in fixed cells

Building upon the solid validation of **6-HoeHESiR** in SMLM of *in vitro* reconstituted nucleosomal arrays and 30-nm chromatin fibers, we then attempted to image native chromatin structures in fixed HeLa cells (Fig. 3a, Supplementary Fig. 7a, Supplementary Video 4). Based on the structural resemblance to *in vitro* reconstituted benchmark chromatin samples, we identified two categories of chromatin structures. The first category consists of nucleosomal arrays (Fig. 3b), which displayed the same pattern of discrete fluorescence signals (Fig. 2d). The second category consists of chromatin fibers with structural plasticity (Fig. 3c, Supplementary Fig. 7b). With the aid of the image analysis program, these fibers were identified with a mean localization precision of 8 nm, average 3D size (FWHM) of 30, 31, 101 nm (x, y, z) and a labeling density of 42,170 molecules/μm^3^ (Fig. 3d, Supplementary Fig. 7d, Supplementary Video 5 and 6). The overlay of widefield images showed that the chromatin fibers reside in both the peripheral and the interior regions of the nuclei (Fig. 3e; Supplementary Fig. 7c). Budding yeast *Saccharomyces cerevisiae* is a unicellular model organism for biological study. Yet, there has been no 3D superresolution imaging of budding yeast chromatin. Therefore, we cultured *S*. *cerevisiae* BY4741 cells and visualized their native chromatin with **6-HoeHESiR** after fixation (Fig. 3f-g, Supplementary Fig. 8a, Supplementary Video 7). We were able to identify nucleosomal arrays and chromatin fibers as the two categories of chromatin structures (Fig. 3h, 3i and 3k, Supplementary Fig. 8b and 8c). Chromatin fibers were visualized with a mean localization precision of 8.4 nm, average 3D size (FWHM) of 34, 34, 113 nm (x, y, z) and a labeling density of 35,403 molecules/μm^3^ (Fig. 3j, Supplementary Fig. 8d, Supplementary Video 8 and 9). Our imaging results provide direct evidence that agrees with recent studies by RICC-seq^27^ and Micro-C genomics^28^ that support the existence of chromatin fibers in mammalian cells and budding yeasts.

**Fig. 3.**
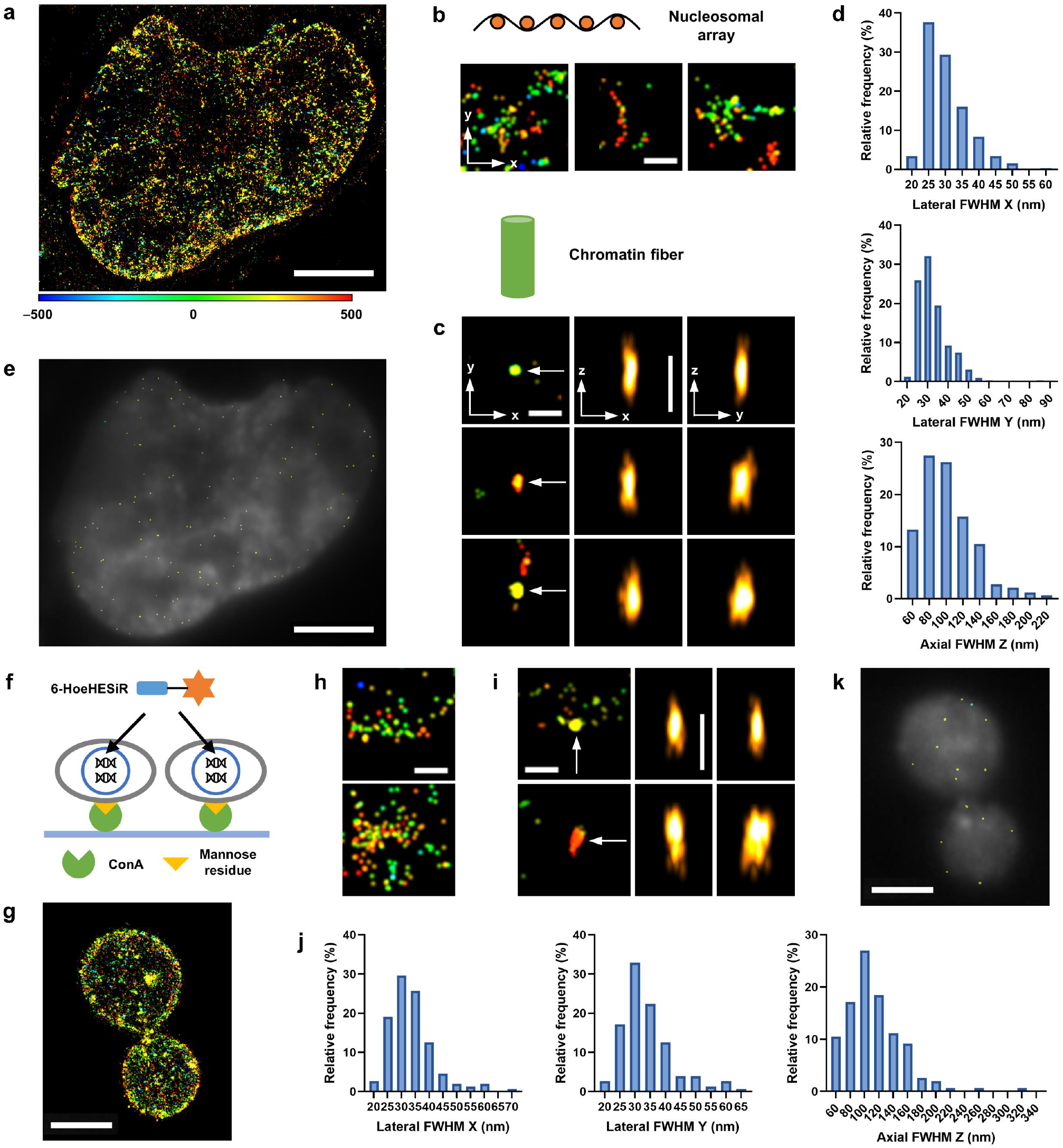
Native chromatin 3D structures in fixed cells. **a,** Image of the nucleus in a HeLa cell. Reconstructed from 5,000 frames (17.7 ms/frame). **b,** Nucleosomal arrays in HeLa cells. **c,** Chromatin fibers in HeLa cells. FWHM (x, y, z) in nm (from top to bottom): 30, 27, 104; 26, 41, 108; 36, 51, 87. Localization precision (nm): 6.9, 6.1, 9.1. **d**, 3D sizes of chromatin fibers by FWHM. *N* = 324 from 3 HeLa cells, identified in 500 frames. **e**, Distribution of chromatin fibers in a HeLa cell. **f,** Immobilization approach for imaging budding yeasts. **g,** The nuclei of two dividing budding yeasts. **h,** Nucleosomal arrays in budding yeasts. **i,** Chromatin fibers in budding yeasts. FWHM (x, y, z) in nm (from upper to lower): 32, 27, 94; 36, 62, 118. Localization precision (nm): 7, 9.2. **d**, 3D sizes of chromatin fibers by FWHM. *N* =152 from 16 budding yeast cells, identified in 500 frames. **k**, Distribution of chromatin fibers in budding yeasts. Scale bar: 5 μm in **a** and **e**; 2 μm in **g** and **k**; 200 nm in **b**, **c**, **h** and **i**. Laser intensity: 3.2 kW/cm^2^ at 656 nm. Axial position (nm) represented by RGB color depth coding.

### Native chromatin 3D structures in living cells

First, we optimized the incubation conditions of **6-HoeHESiR** with living HeLa cells by light-sheet microscopy (Fig. 4a-b, Supplementary Fig. 9a-b). To our delight, **6-HoeHESiR** exhibited good cell permeability and high selectivity to provide 3D visualization of chromatin structures in whole nuclei. Without the Hoechst fragment, **6-HESiR** did not stain any nuclear component (Supplementary Fig. 9c). The cytotoxicity assay showed that **6-HoeHESiR** caused negligible toxicity with a short incubation time (Supplementary Fig. 9d). Subsequent 3D SMLM experiments showed that **6-HoeHESiR** maintained its excellent self-blinking capacity in living HeLa cells (Supplementary Video 10), and native chromatin structures were clearly revealed (Fig. 4c; Supplementary Fig. 10a). We identified two categories of chromatin structures in living HeLa cells, namely, nucleosomal arrays and chromatin fibers (Fig. 4d, 4e and 4g, Supplementary Fig. 10b and 10d). Using the image analysis program, these fibers were identified with a mean localization precision of 7.4 nm, average 3D size (FWHM) of 36, 39, 103 nm (x, y, z) and a labeling density of 32,149 molecules/μm^3^ (Fig. 4f, Supplementary Fig. 10c, Supplementary Videos 11 and 12). The statistical distribution of 3D sizes and long-to-short fiber diameter ratios and the heterogeneity of visualized morphologies reflect the structural polymorphism of chromatin fibers. Our 2D imaging results are consistent with recent reports^29,30,23^ that nucleosome clutches or chromatin domains were observed with no clear evidence of horizontally or randomly oriented chromatin fibers. However, our 3D analysis, performed both manually and by the computer analysis program, identified chromatin fibers oriented axially. We propose that such an ordered chromatin fiber orientation might not only be consistent with highly regulated replication and transcription processes but facilitate these processes. Remarkably, native chromatin structures in the nucleus are quite dynamic (Supplementary Video 13). We captured transient chromatin fibers with a temporal resolution of less than 2 seconds (Fig. 4h; Supplementary Video 14). The mean resting/residing lifetime of chromatin fibers was estimated to be 0.6 seconds (Supplementary Fig. 10c). This dwelling lifetime is comparable to the time scale of folding and unfolding dynamics of *in vitro*-reconstituted 30-nm chromatin fibers revealed by single-molecule force spectroscopy^26^. When the laser intensity was reduced to a modest level of 950 W/cm^2^ from 3.2 kW/cm^2^ to alleviate phototoxicity, nucleosomal arrays were visualized again, and we could still identify chromatin fibers at sufficient resolution with a mean localization precision of 8.8 nm, average 3D size (FWHM) of 35, 39, 108 nm (x, y, z) and a labeling density of 29,691 molecules/μm^3^ (Supplementary Fig. 11). Fast chromatin dynamics were observed (Supplementary Video 15), and the mean resting lifetime of chromatin fibers was estimated to be 0.6 seconds, consistent with the result obtained at 3.2 kW/cm^2^ laser intensity.

**Fig. 4.**
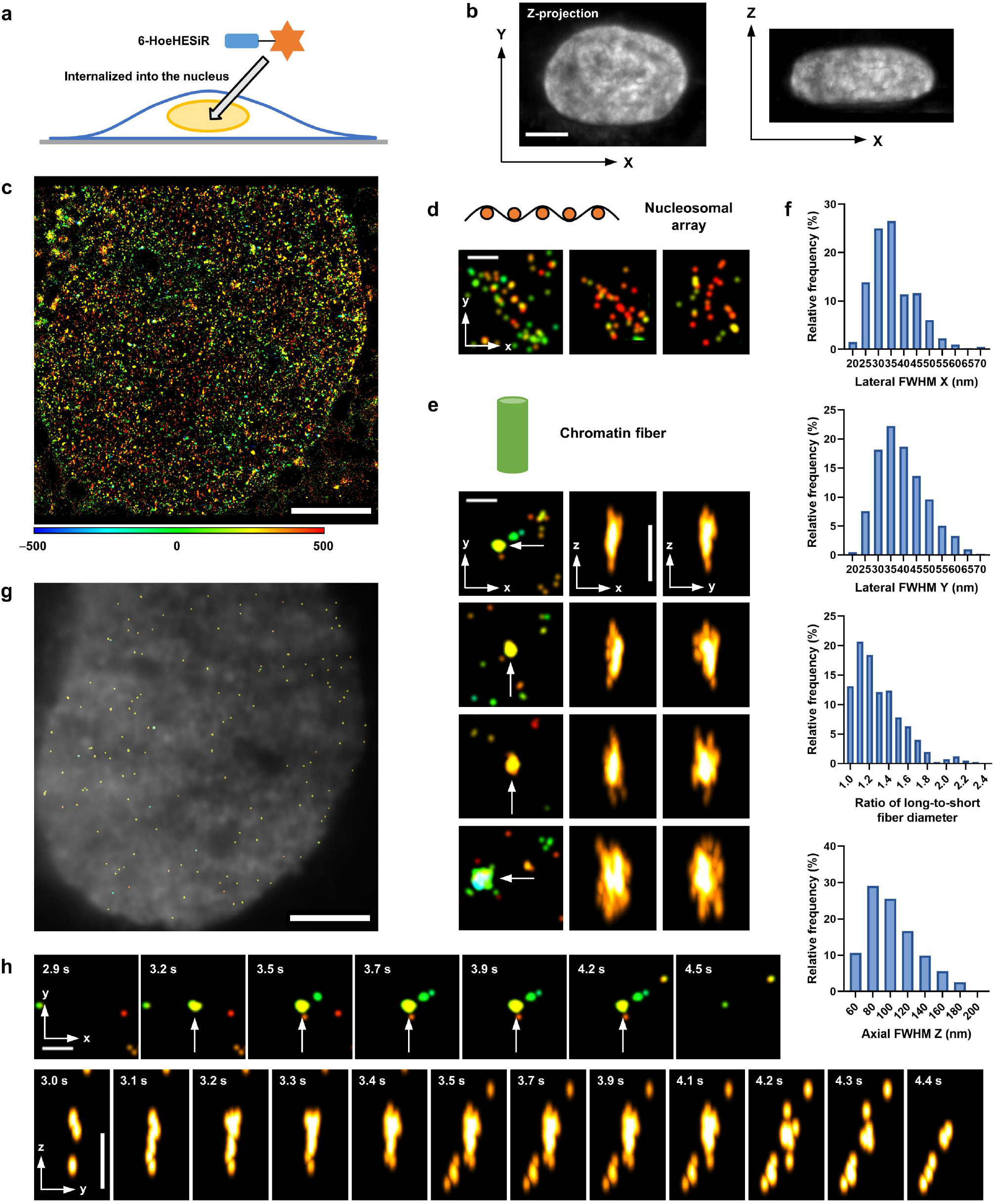
Native chromatin 3D structures in living HeLa cells. **a,** Schematic illustration of chromatin labeling with **6-HoeHESiR** (5 μM for 30 min). **b,** Images of a whole nucleus by light-sheet microscopy. **c**, Image of a nucleus. Reconstructed with 2,000 frames (17.7 ms/frame). **d**, Nucleosomal arrays. **e**, Chromatin fibers. FWHM (x, y, z) in nm (from top to bottom): 30, 30, 107; 35, 42, 124; 34, 51, 102; 69, 51, 154. Localization precision (nm): 7.6, 6.8, 6.8, 9.5. **f**, 3D sizes of chromatin fibers by FWHM. *N* = 396 from 4 cells, identified in 500 frames. **g**, Distribution of chromatin fibers (identified in 500 frames). **h**, Fast dynamics of a chromatin fiber. Snapshots from Supplementary Videos 7 and 8 (Supplementary Information, video legends). Scale bar: 5 μm in **b**; **c** and **g**; 200 nm in **d**, **e** and **h**. Laser intensity: 3.2 kW/cm^2^ at 656 nm. Axial position (nm) represented by RGB color depth coding.

A previous cryo-electron tomography (cryo-ET) study identified short chromatin fibers with 30 nm in diameter in isolated nuclei from chicken erythrocytes^31^. Therefore, we also attempted 3D SMLM experiments on living chicken erythrocytes with **6-HoeHESiR**. By applying a modest laser intensity of 950 W/cm^2^, 3D chromatin structures in chicken erythrocyte nuclei were successfully visualized (Fig. 5a-c; Supplementary Fig. 12a; Supplementary Video 16). The same two categories of chromatin structures were observed, and chromatin fibers were identified with a mean localization precision of 8 nm, average 3D size (FWHM) of 32, 35, 90 nm (x, y, z) and a higher labeling density of 56,494 molecules/μm^3^ (Fig. 5d-g; Supplementary Fig. 12b-d; Supplementary Videos 17 and 18). The fast dynamics of chromatin structures in chicken erythrocyte nuclei were also observed (Supplementary Video 19), with the mean resting lifetime of chromatin fibers estimated to be 1 second.

**Fig. 5.**
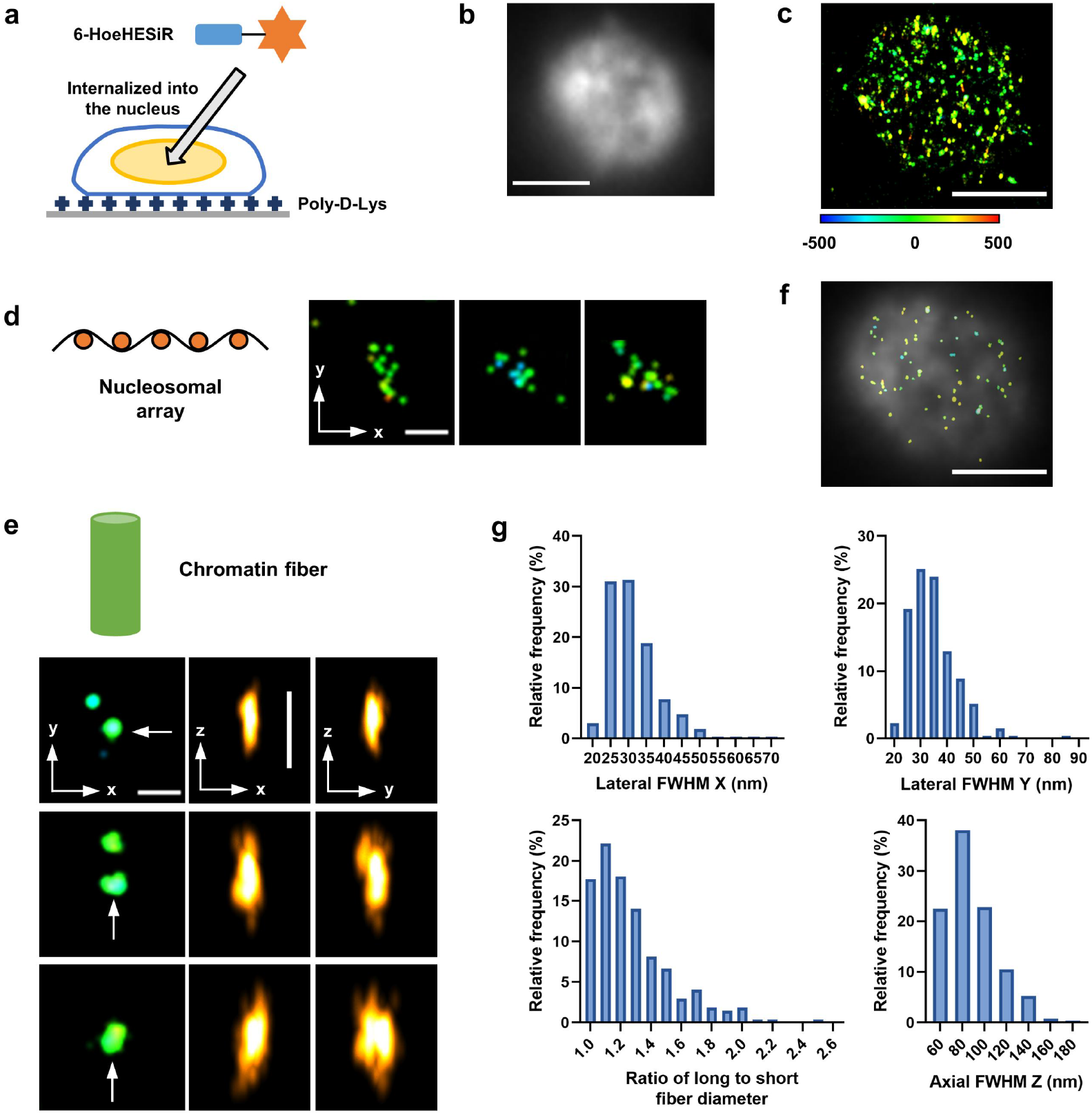
Native chromatin 3D structures in living chicken erythrocytes. **a,** Schematic illustration of chromatin labeling with **6-HoeHESiR** (0.5 μM for 10 min). **b,** Image of a nucleus by widefield microscopy. **c**, Image of a nucleus. Reconstructed with 10,000 frames (17.7 ms/frame). **d**, Nucleosomal arrays. **e**, Chromatin fibers. FWHM (x, y, z) in nm (from top to bottom): 27, 29, 83; 37, 40, 124; 40, 62, 95. Localization precision (nm): 6.8, 6.9, 8.1. **f**, 3D sizes of chromatin fibers by FWHM. *N* = 268 from 3 cells, identified in 5,000 frames. **g**, Distribution of chromatin fibers (identified in 5,000 frames). Scale bar: 2 μm in **b**, **c** and **g**; 200 nm in **d** and **e**. Laser intensity: 0.95 kW/cm^2^ at 656 nm. Axial position (nm) represented by RGB color depth coding.

### The connection between 3D chromatin organization and histone acetylation

It is known that histone acetylation is a critical epigenetic marker for chromatin fiber relaxation and optimal transcription, whereas histone deacetylation is required for chromatin fiber condensation and transcriptional repression^32^. Inhibition of histone deacetylases (HDACs), key enzymes that reduce histone acetylation, is an important approach to anticancer drug development by inducing chromatin decompaction and reactivating tumor suppressor genes^33,34^. To investigate the connection between chromatin fibers and histone acetylation status in living cells, we selected three HDAC inhibitors (Fig. 6a): 1) trichostatin A (TSA), a widely used HDAC pan-inhibitor; 2) entinostat, a class I HDAC 1 – 3 selective inhibitor; and 3) ricolinostat, a class IIb HDAC 6 selective inhibitor. Live HeLa cells were treated with each of these HDAC inhibitors and then imaged by 3D SMLM (Fig. 6b). The image analysis results show that the density of chromatin fibers decreased by 52% and 63% upon treatments with TSA and entinostat, respectively^35,36^, whereas no significant change in fiber density was observed upon ricolinostat treatment (Fig. 6c-d, Supplementary Fig. 13-15). These results are consistent with literature reports that the functions of class I HDACs 1–3 are exclusively restricted to the nucleus, while class IIb HDAC 6 functions largely outside the nucleus by mediating the deacetylation of cytosolic proteins^37^. Taken together, our live-cell 3D imaging results provide the first live-cell evidence for the global change in 3D chromatin organization in response to histone hyperacetylation induced by small-molecule HDAC inhibitors.

**Fig. 6.**
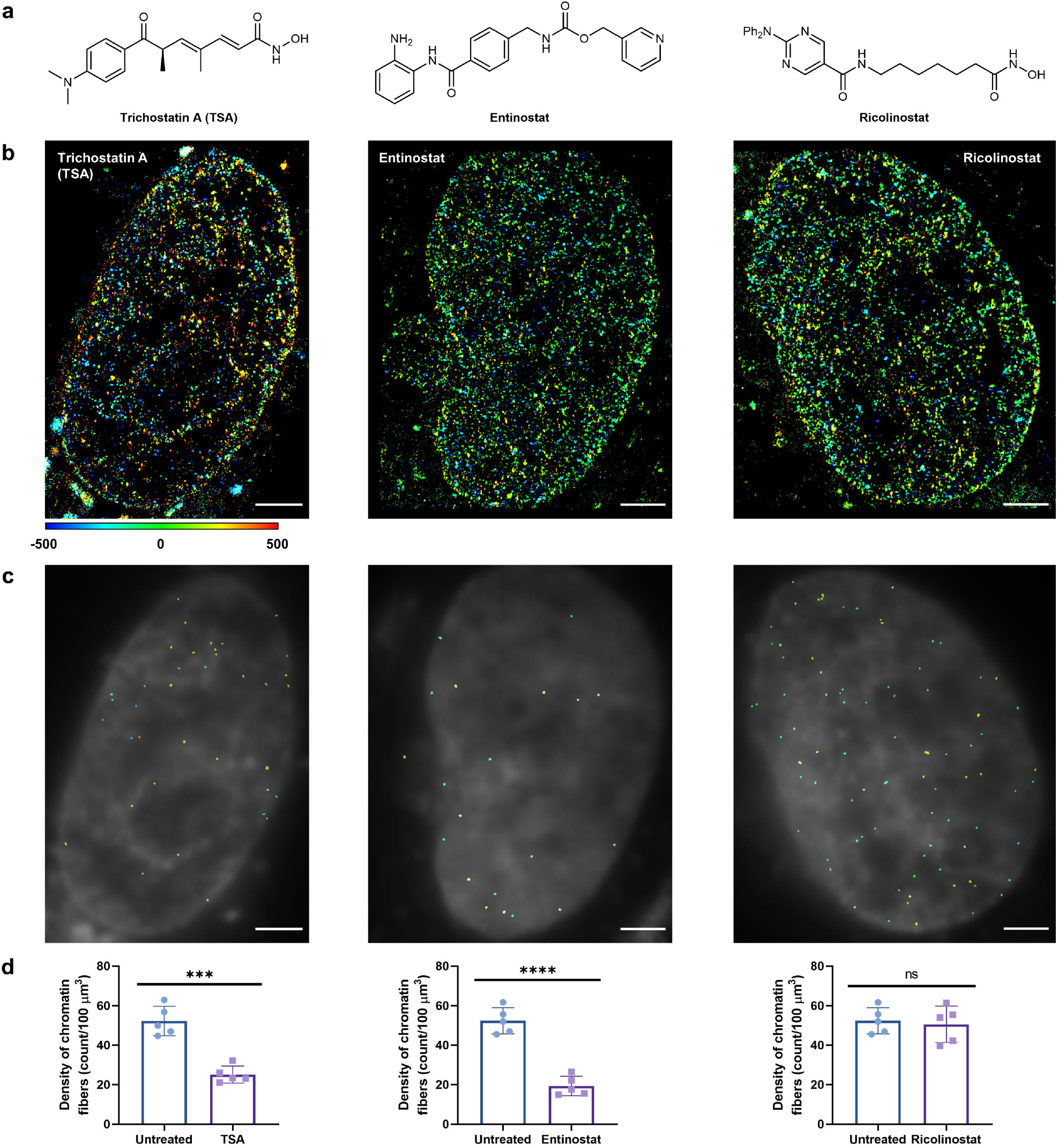
Decompaction of chromatin fibers induced by HDAC inhibitors. **a**, Structures of small-molecule inhibitors of HDAC. **b**, Images of the nuclei in treated HeLa cells. Reconstructed with 2,000 frames (17.7 ms/frame). **c**, Distribution of chromatin fibers (identified in 500 frames). **d**, Density of chromatin fibers (identified in 500 frames). Data are individual values and mean ± s.d., *N* =5. Statistical significance was determined by *t test* with *P* = 0.0001, <0.0001 and 0.7278 (from left to right). The scale bar is 2 μm. Laser intensity: 3.2 kW/cm^2^ at 656 nm. Axial position (nm) represented by RGB color depth coding. Cells were treated with TSA (200 nM), entinostat (2 μM), and ricolinostat (1 μM) for 20 h, respectively, and then incubated with **6-HoeHESiR** (5 μM) for 30 min.

## DISCUSSION

To summarize, we have demonstrated the success of developing a self-blinking Si-rhodamine dye **6-HESiR** and a self-blinking fluorogenic DNA probe **6-HoeHESiR** by structurally fine-tuning the “ON” state population to a suitable level. This design strategy has proven effective in 3D imaging of *in vitro* reconstituted chromatin samples and native chromatin in both fixed cells and living cells by SMLM. It is noteworthy that high spatiotemporal resolution and labeling density were achieved in live-cell 3D SMLM even at a reduced laser intensity that alleviates phototoxicity. Consequently, we were able to observe native chromatin organized in hierarchical forms of nucleosomal arrays and chromatin fibers with fast dynamics and structural polymorphism. Our imaging results are consistent with recent RICC-seq^27^ and high-resolution Micro-C^28^ studies that support the existence of chromatin fibers in mammalian cells and budding yeasts. To the best of our knowledge, we have obtained the first live-cell evidence for a hierarchical model of chromatin organization^38–40^ (Fig. 7), through which the 2-m long genomic DNA is highly compacted and packaged into the 10-μm eukaryotic nucleus. The structural plasticity/heterogeneity and fast dynamics of chromatin fibers are consistent with the complex regulatory environment in the nucleus, such as histone modifications and variants, nonhistone architectural proteins, and linker DNA lengths for various biological functions^38–40^. The density/population of chromatin fibers was reduced by small-molecule inhibitors of HDACs that are responsible for histone deacetylation. For the first time, this phenomenon provides a live-cell 3D view of the tight connection between the histone acetylation state and chromatin hierarchical organization with unprecedented resolution. In light of our initial success, we anticipate further in-depth studies on the structures and functions of chromatin fibers in DNA-related processes in living cells. We also envision that our structural fine-tuning strategy could be implemented to develop more self-blinking fluorophores and probes to enable the study of subcellular organelles and biological events by live-cell 3D superresolution imaging.

**Fig. 7.**
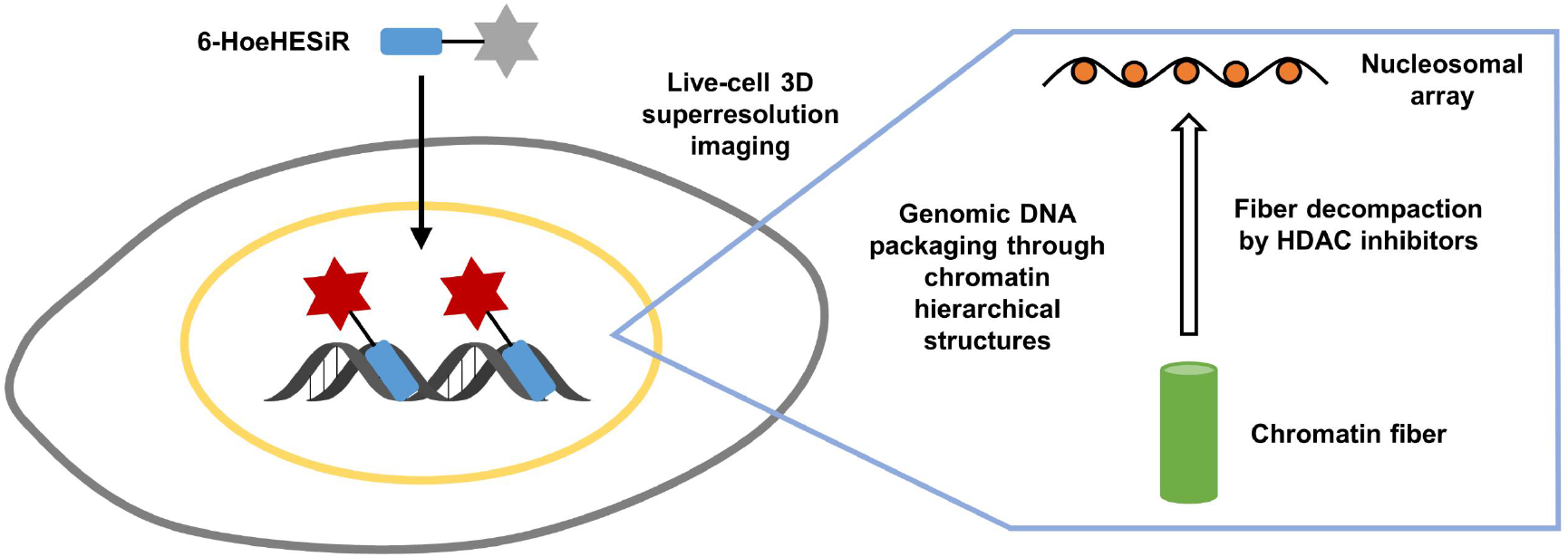
Schematic illustration of genomic DNA packaging via hierarchical chromatin structures.

## Acknowledgments

We thank the HKU Li Ka Shing Faculty of Medicine Imaging and the Flow Cytometry Core of CPOS for support in cell imaging. This work was supported by The University of Hong Kong, Westlake University, Morningside Foundation, and Hong Kong Research Grants Council under the Area of Excellence Scheme (AoE/P-705/16 to D.Y.). This work was supported by grants for G.L. from the National Natural Science Foundation of China (31991161), the Ministry of Science and Technology of China (2017YFA0504202), the Beijing Municipal Science and Technology Commission (Z201100005320013) and the HHMI International Research Scholars Program (55008737). R. L. was supported by the ECS (grant number 27204518) of the Hong Kong SAR government and by the URC fund at the University of Hong Kong. We thank Dr. Teng Zhao and Dr. Zhe Hu for their technical support in light-sheet microscopy. We thank Dr. Zhe Hu, Dr. Teng Zhao, Dr. Shengwang Du and Dr. Wei Li for helpful discussions. We thank Dr. Nai-Kei Wong and Ms. Jasmine Chit Ying Lau for proofreading the manuscript.

## Author contributions

Y.Z. and G.L. conceived the idea. G.L. and D.Y. supervised the project. Y.Z. designed **6-HESiR** and **6-HoeHESiR**. Y.Z. performed chemical synthesis and characterization with contribution from R.X. Y.Z. studied photophysical properties with contribution from S.Y. C.L. prepared *in vitro* reconstituted samples and performed electron microscopy analysis. S.-H.L. performed the SMLM experiment with lambda DNA and analyzed the data. Y.Z., S.Y. and S.D. performed cell culture and treatment. Z.Z. performed budding yeast culture. Y.Z. and S.Y. performed SMLM experiments with reconstituted chromatin structures and with fixed and live cells. Y.C. and Y.L. performed the SMLM validation experiments. L.H. performed cytotoxicity assay. S.-M.L. developed the computer program and the web server for chromatin fiber analysis under the supervision of R.L. Y.Z. analyzed chromatin imaging data and performed statistical analysis with contribution from S.Y. and S.-M.L. Y.Z., S.Y., G.L. and D.Y. interpreted imaging results with contribution from all other authors. Y.Z., S.Y., G.L. and D.Y. wrote the manuscript with input from all other authors.

## Competing interests

The authors declare no competing interests.

## Data availability

All data that support the findings of this manuscript are provided in text, figures, and supplementary videos. Original data are provided upon reasonable request to the correspondence authors.

## Notes

### Competing Interest Statement

The authors have declared no competing interest.

### Summary of Updates

Added fixed-cell results. Revised text, figures and videos

