## Supplementary Information for "A self-blinking DNA probe for 3D superresolution imaging of native chromatin"

#### Supplementary figures

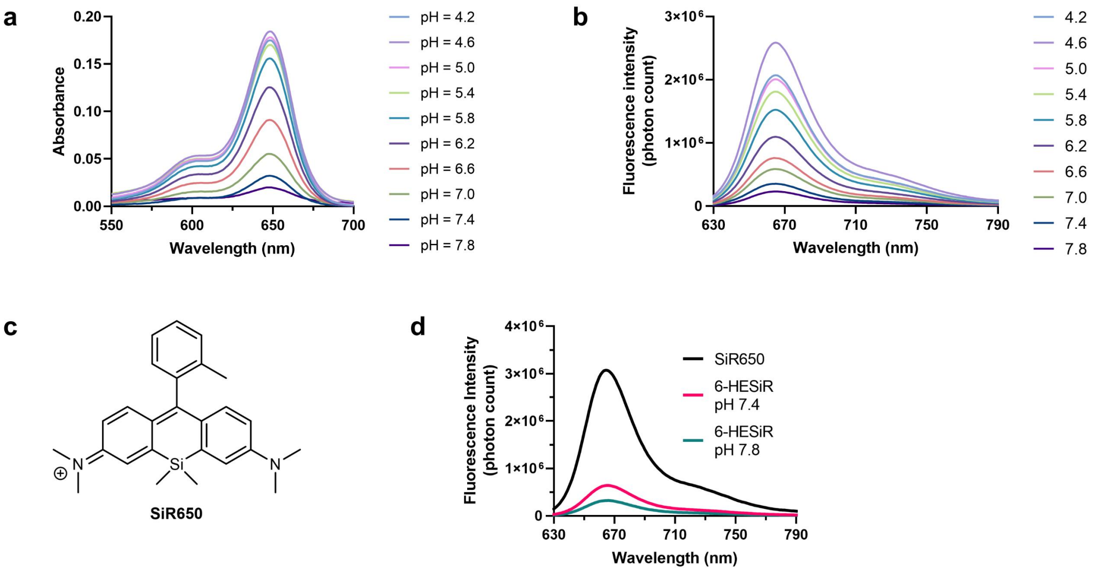


**Supplementary Fig. 1 Photophysical properties of 6-HESiR. a**, Absorption spectra of **6-HESiR** (2 µM) at various pH values in 0.1 M potassium phosphate buffer. **b**, Fluorescence emission spectra of **6-HESiR** (0.5 µM) at various pH values in 0.1 M potassium phosphate buffer. **c**, Structure of **SiR650**. **d**, Fluorescence emission spectra of **6-HESiR** (2 µM) in 0.1 M potassium phosphate buffer (pH = 7.4 and 7.8) with **SiR650** (2 µM) as the "100%" benchmark.

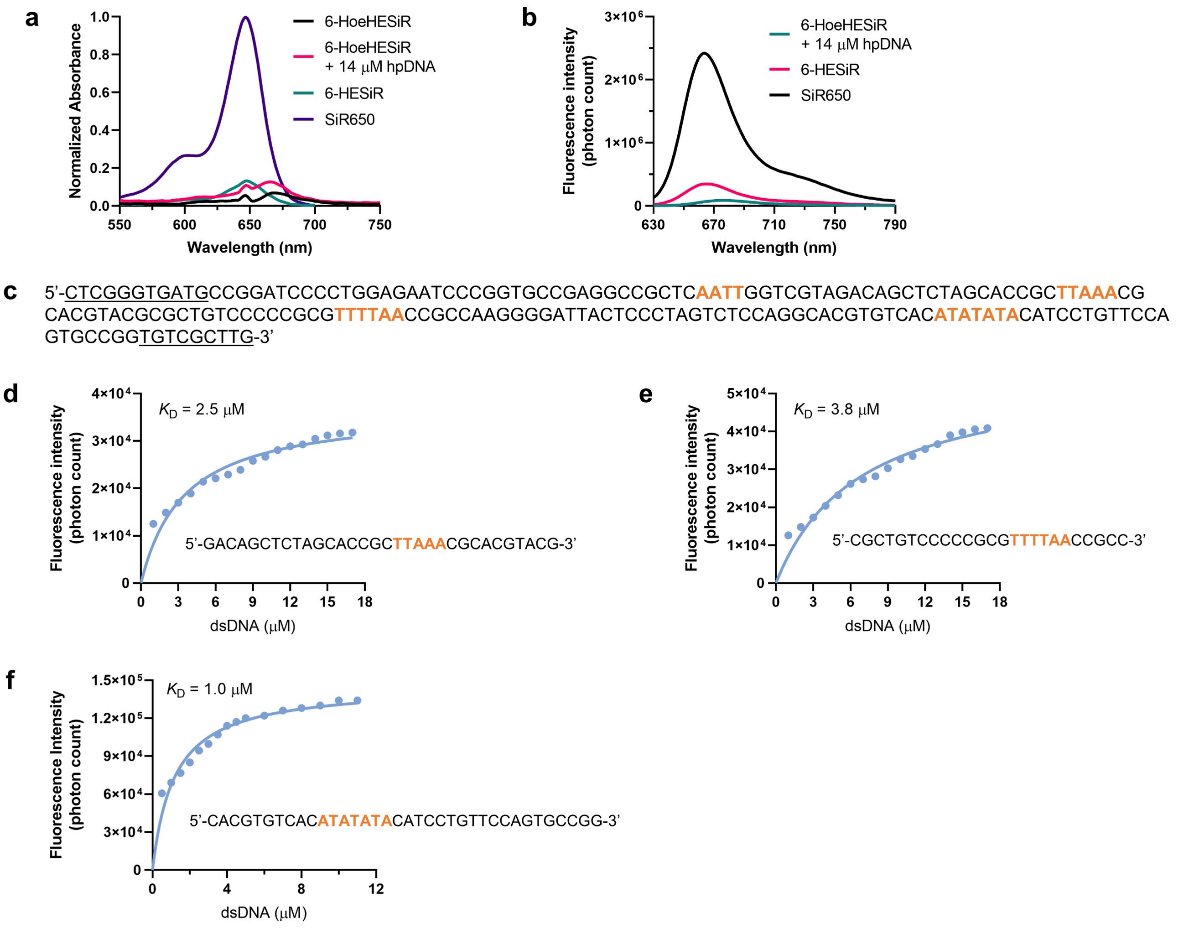


**Supplementary Fig. 2 Fluorescence titration of 6-HoeHESiR. a**, Absorbance of **6-HoeHESiR** (0.5 μM) in Tris-HCl saline buffer (50 mM Tris-HCl, 100 mM NaCl, pH = 7.4). **b**, Fluorescence of **6-HoeHESiR**, **6-HESiR** and **SiR650** (0.5 μM) in Tris-HCl saline buffer. **c**, Sequence of the 187-bp DNA template for reconstituted chromatin samples. The linker DNA sequence is underlined. **d-f**, Fluorescence titration with 0.5 μM **6-HoeHESiR** and dsDNA segments in the Tris-HCl saline buffer.

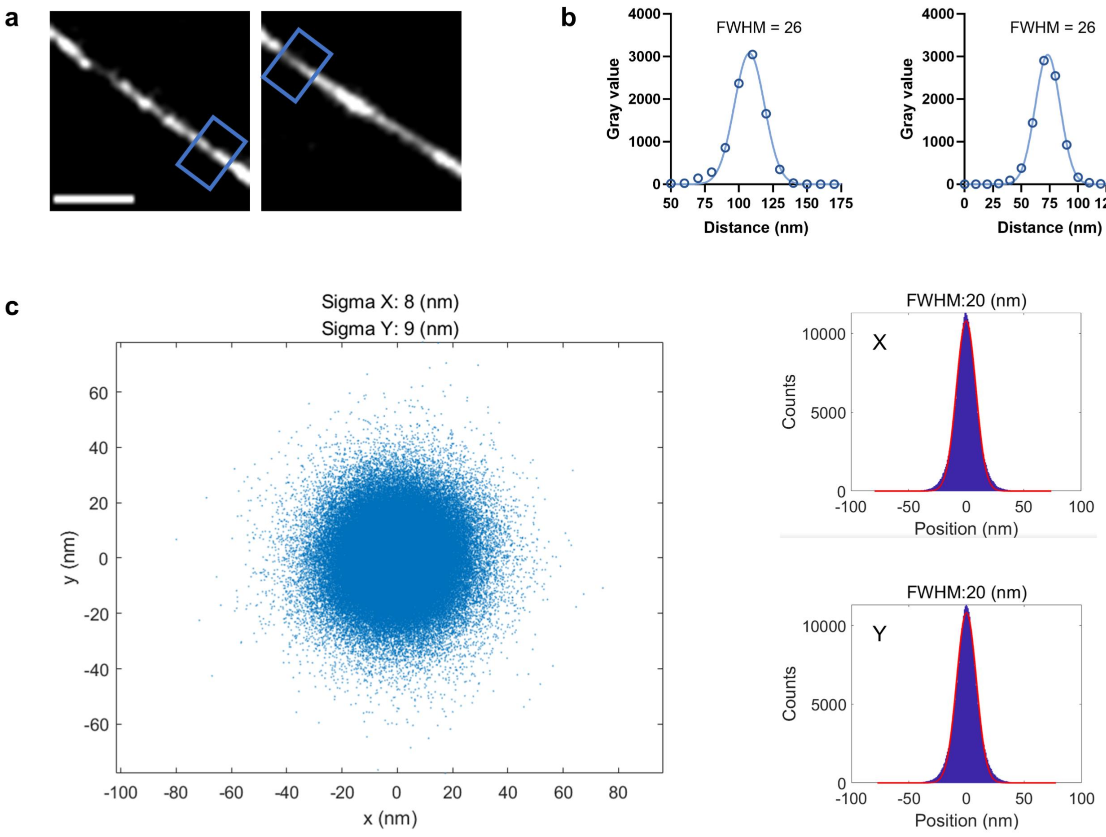


**Supplementary Fig. 3 SMLM with lambda DNA. a**, Reconstructed images of spin-coated λ-DNA. **b**, Line profiles for the boxed regions in **a** for obtaining FWHM values. **c**, Localization precision. Imaging performed with 100 nM **6-HoeHESiR**. Scale bar: 200 nm.

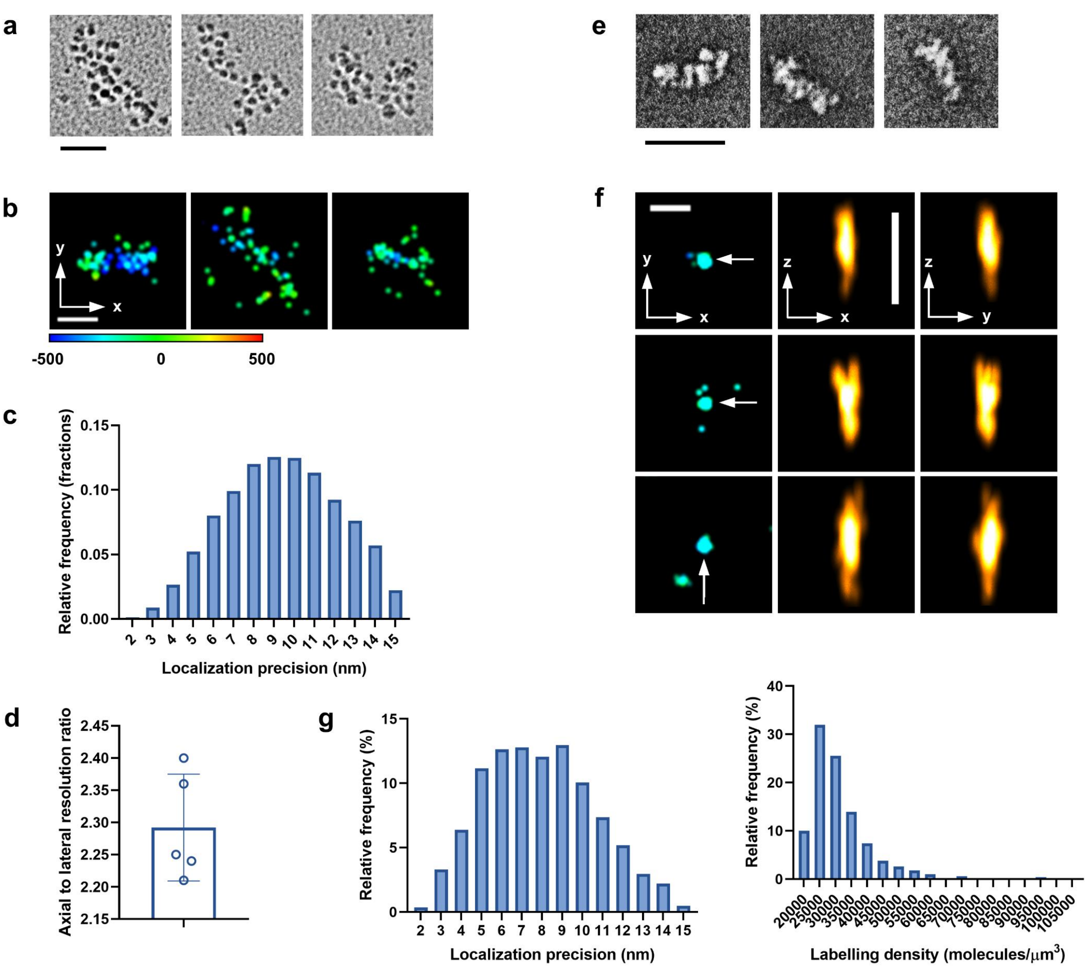


**Supplementary Fig. 4 SMLM with *in vitro* reconstituted chromatin structures. a**, EM images of 40×187-bp nucleosomal arrays (by metal-shadowing). **b**, Images of 40×187-bp nucleosomal arrays by 3D SMLM. **c**, Localization precision for 40×187-bp nucleosomal arrays. **d**, Ratio of axial resolution to lateral resolution. Data are mean ± s.d., $N = 5$. **e**, EM images of 40×187-bp 30-nm chromatin fibers (by negative staining). **f**, Images of 40×187-bp 30-nm fibers by 3D SMLM. FWHM (x, y, z) in nm (from top to bottom): 26, 30, 93; 27, 29, 96; 29, 31, 99. Localization precision (nm): 6.2, 6.8, 6.3. **g**, Localization precision and labeling density for identified 30-nm chromatin fibers. Incubation with **6-HoeHESiR** (2.5 μM) for 10 min. Scale bar: 100 nm in **a** and **e**; 200 nm in **b** and **f**. Laser intensity: 0.95 $kW/cm^2$ at 656 nm. Axial position (nm) represented by RGB color depth coding.

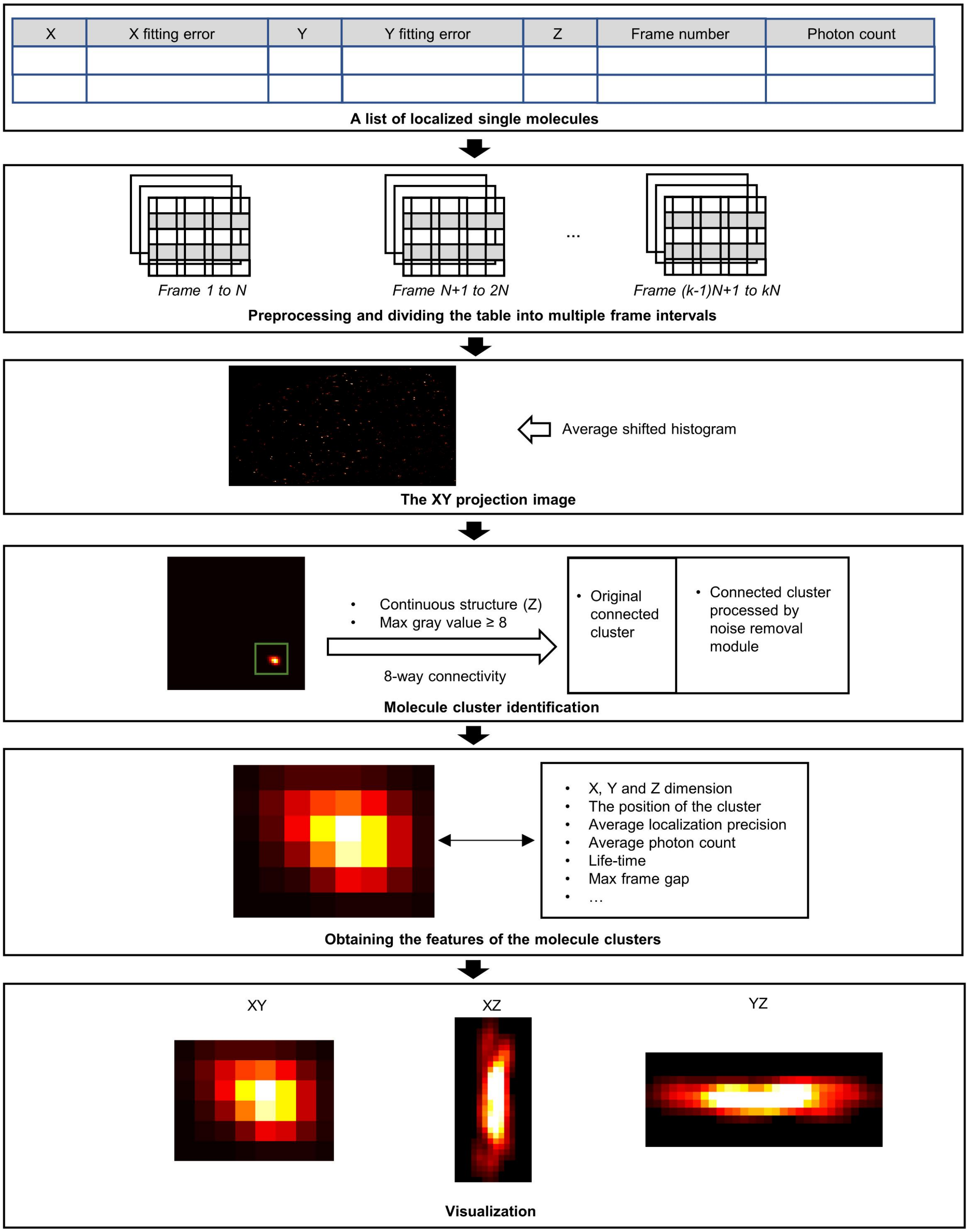


**Supplementary Fig. 5** The flowchart of the analysis of molecular clusters for identification of chromatin fibers.

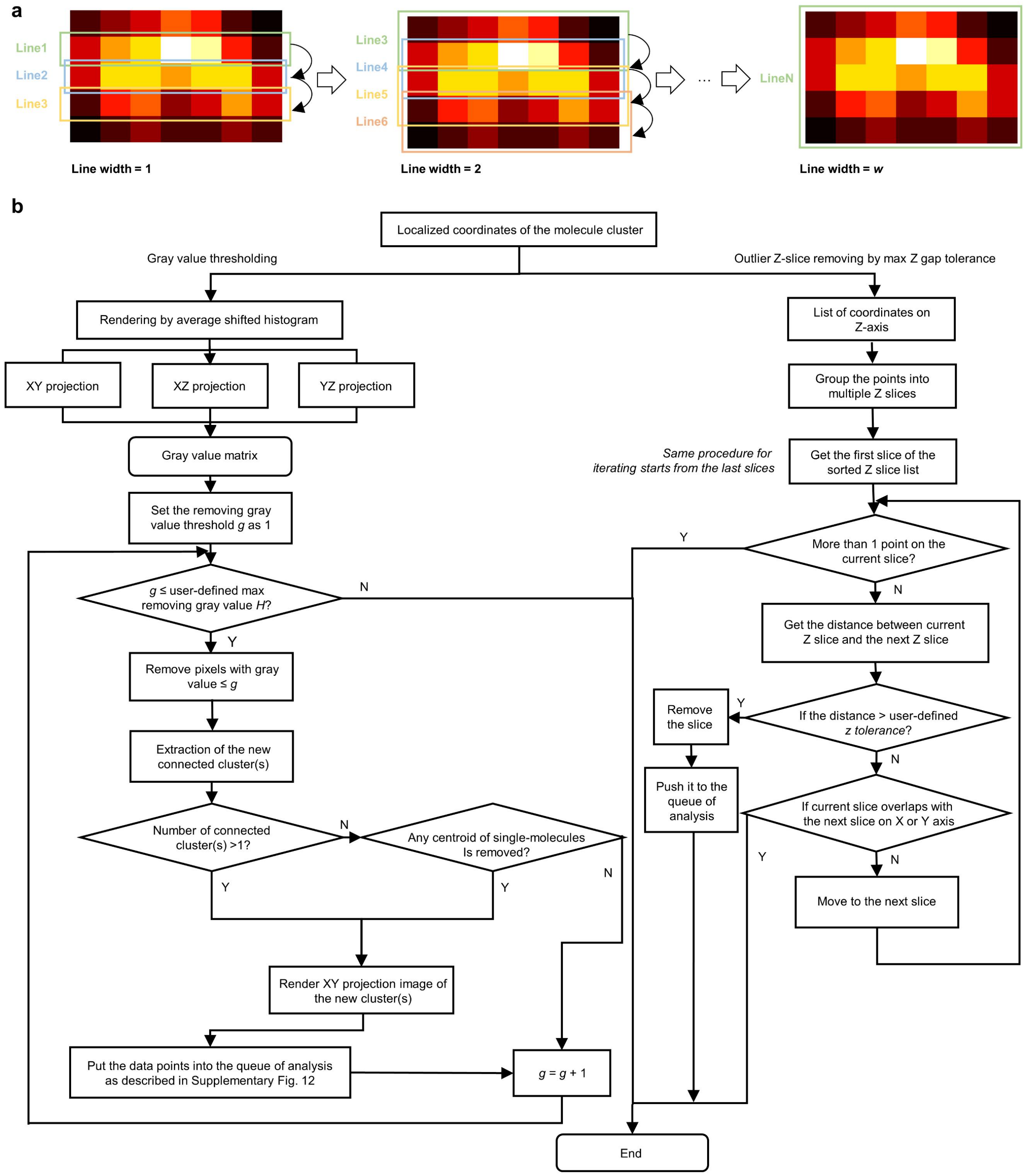


**Supplementary Fig. 6 Chromatin fiber analysis program. a**, The demonstration of the moving line strategy for estimating X dimension of molecular clusters. **b**, The flowchart of the noise removal module, which consists of two submodules: gray value thresholding and outlier Z slice removing by max Z gap tolerance. *g* ranges from 1 to the user-defined max threshold *H*.

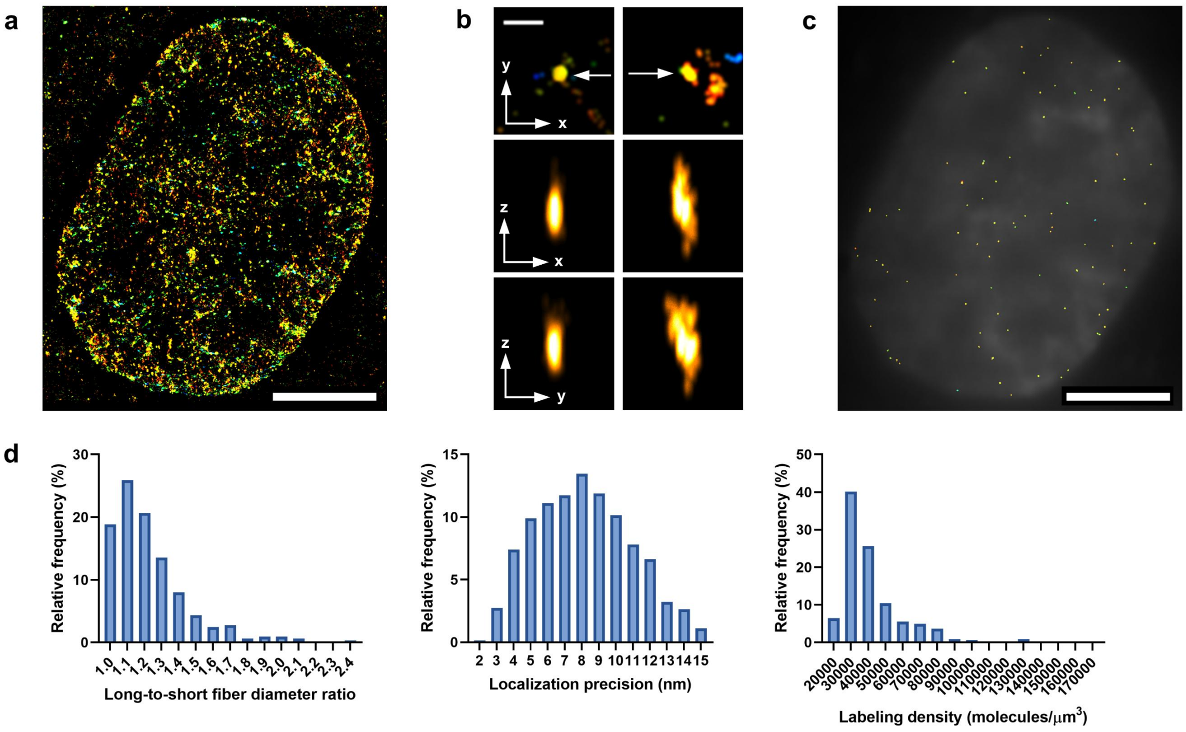


**Supplementary Fig. 7 Chromatin fibers in fixed HeLa cells. a**, The image of a nucleus. Reconstructed from 5,000 frames (17.7 ms/frame). **b**, Chromatin fibers. FWHM (x, y, z) in nm (from left to right): 28, 30, 84; 32, 45, 114. Localization precision (nm): 6.1, 6.6. **c**, The distribution of chromatin fibers (identified in 500 frames). **d**, Long-to-short fiber diameter ratio, localization precision and labeling density for identified fibers. Incubation with **6-HoeHESiR** (8 μM) for 15 min. Scale bar: 5 μm in **a** and **c**; 200 nm in **b**. Laser intensity: 3.2 kW/cm$^2$ at 656 nm. Axial position (nm) represented by RGB color depth coding.

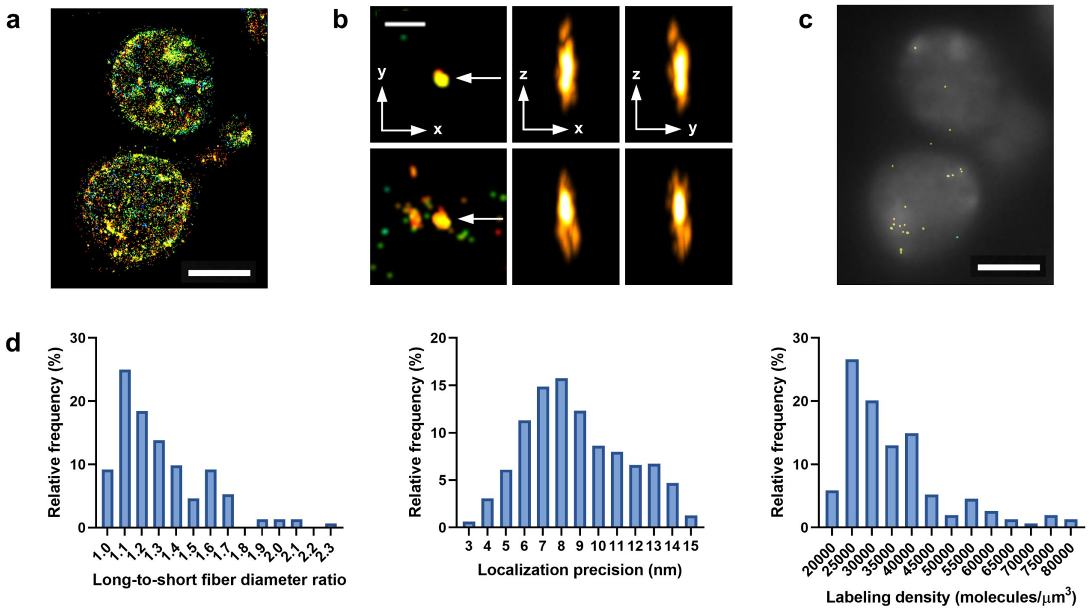


**Supplementary Fig. 8 Chromatin fibers in budding yeasts. a**, The image of a nucleus. Reconstructed from 5,000 frames (17.7 ms/frame). **b**, Chromatin fibers. FWHM (x, y, z) in nm (from upper to lower): 31, 29, 124; 38, 32, 98. Localization precision (nm): 7.3, 7.1. **c**, The distribution of chromatin fibers (identified in 500 frames). **d**, Long-to-short fiber diameter ratio, localization precision and labeling density for identified fibers. Incubation with **6-HoeHESiR** (8 μM) for 15 min. Scale bar: 2 μm in **a** and **c**; 200 nm in **b**. Laser intensity: 3.2 kW/cm$^2$ at 656 nm. Axial position (nm) represented by RGB color depth coding.

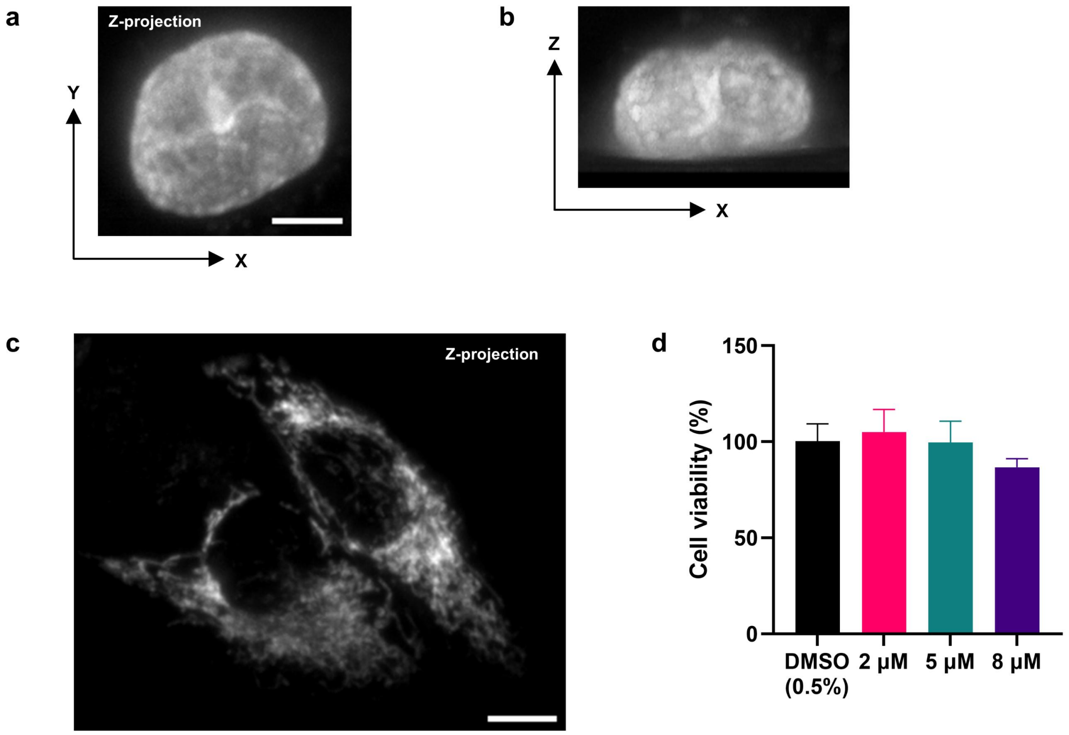


**Supplementary Fig. 9 Light-sheet microscopy of living HeLa cells. a**, The Z-projection image of the nucleus in a living HeLa cell. **b,** The XZ projection image of the same nucleus. **c**, The Z-projection image of living HeLa cells stained with **6-HESiR**. **d**, Cytotoxicity assay with **6-HoeHESiR** (5 h incubation in DMEM). $N = 3$, data are mean + SD. Incubation with **6-HoeHESiR** (**a** and **b**, 5 µM) and **6-HESiR** (**c**, 2 µM) for 30 min, respectively. Scale bar: 5 µm in **a** and **b**; 10 µm in **c**.

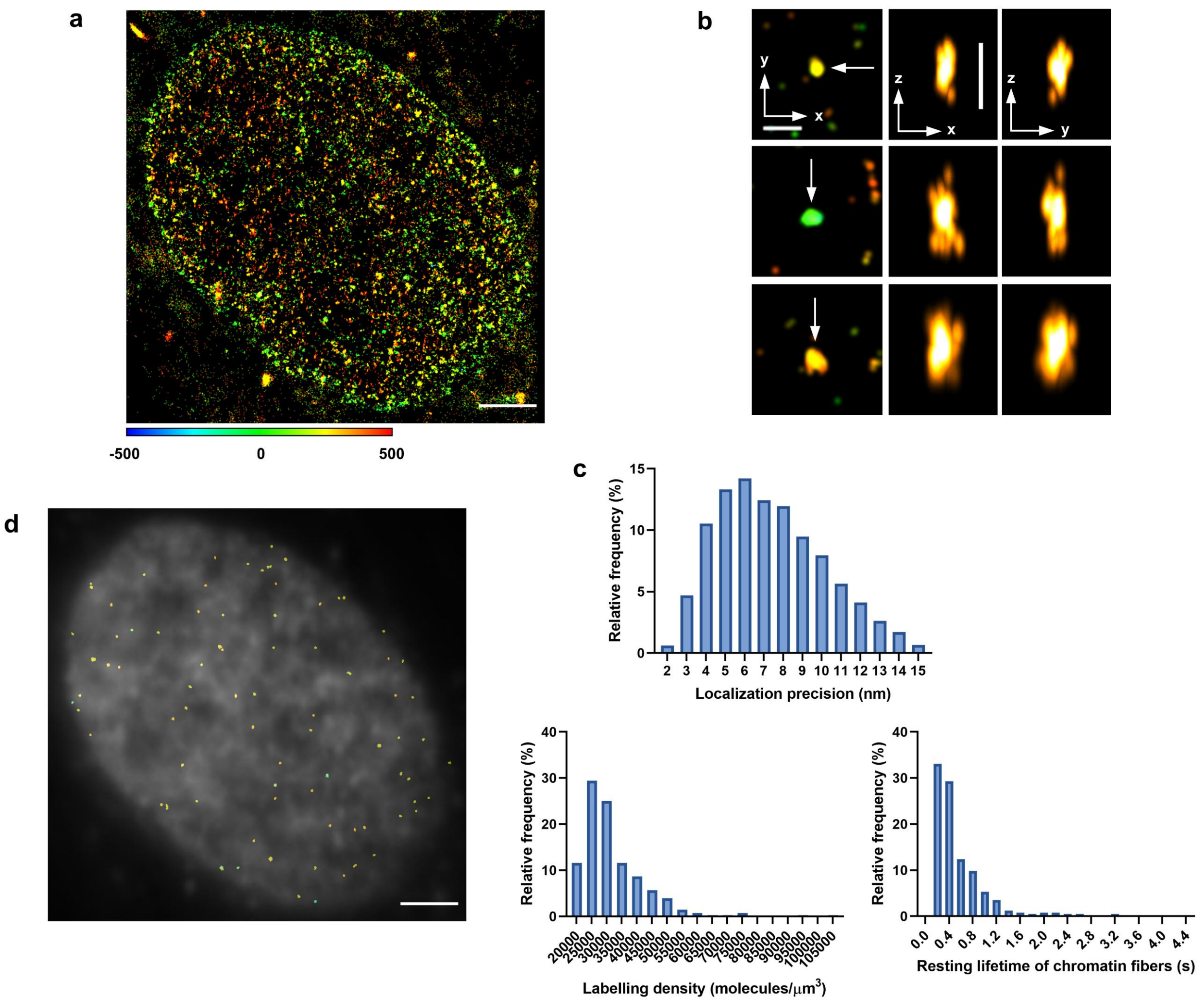


**Supplementary Fig. 10 Chromatin fibers in living HeLa cells. a**, The image of a nucleus. Reconstructed from 2,000 frames (17.7 ms/frame). **b**, Chromatin fibers. FWHM (x, y, z) in nm (from top to bottom): 31, 34, 94; 44, 35, 116; 45, 50, 103. Localization precision (nm): 7, 7.2, 9.1. **c**, Localization precision, labeling density, and resting lifetime for identified fibers in 500 frames. **d**, The distribution of chromatin fibers (identified in 500 frames). Incubation with **6-HoeHESiR** (5 μM) for 30 min. Scale bar: 2 μm in **a** and **d**; 200 nm in **b**. Laser intensity: 3.2 kW/cm$^2$ at 656 nm. Axial position (nm) represented by RGB color depth coding.

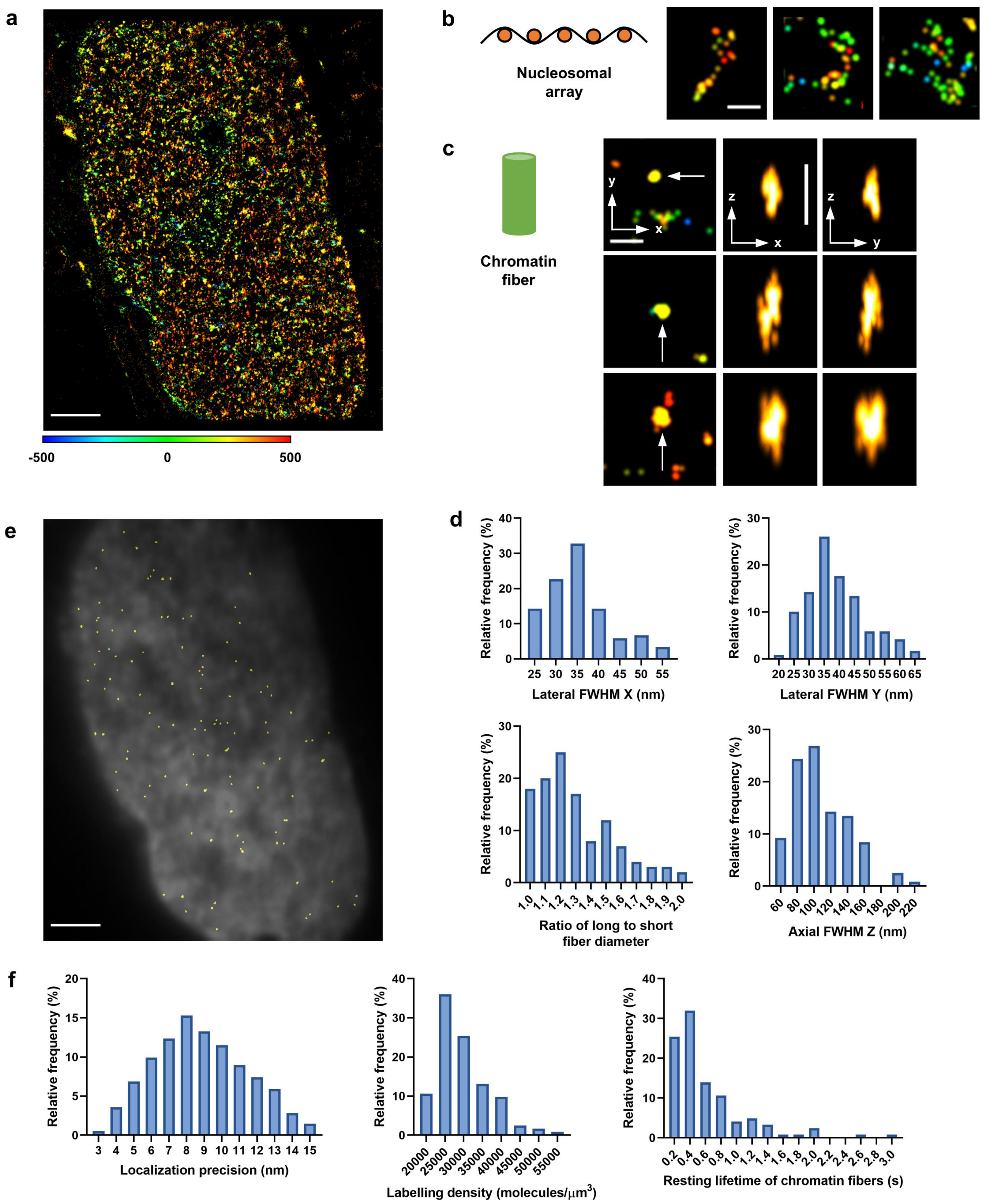


**Supplementary Fig. 11 Native chromatin 3D structures revealed with reduced phototoxicity. a**, The image of the nucleus in a live HeLa cell. Reconstructed from 2,000 frames (17.7 ms/frame). **b**, Images of nucleosomal arrays. **c**, Chromatin fibers. FWHM (x, y, z) in nm (from top to bottom): 31, 33, 95; 42, 36, 130; 47, 62, 112. Localization precision (nm): 6.8, 7.2, 8.9. **d**, 3D sizes of identified fibers in 500 frames. $N = 119$. **e**, The distribution of chromatin fibers (identified in 500 frames). **f**, Localization precision, labeling density, and resting lifetime for identified fibers in 500 frames. Incubation with **6-HoeHESiR** (5 µM) for 30 min. Scale bar: 2 µm in **a** and **e**; 200 nm in **b** and **c**. Laser intensity: 0.95 kW/cm$^2$ at 656 nm. Axial position (nm) represented by RGB color depth coding.

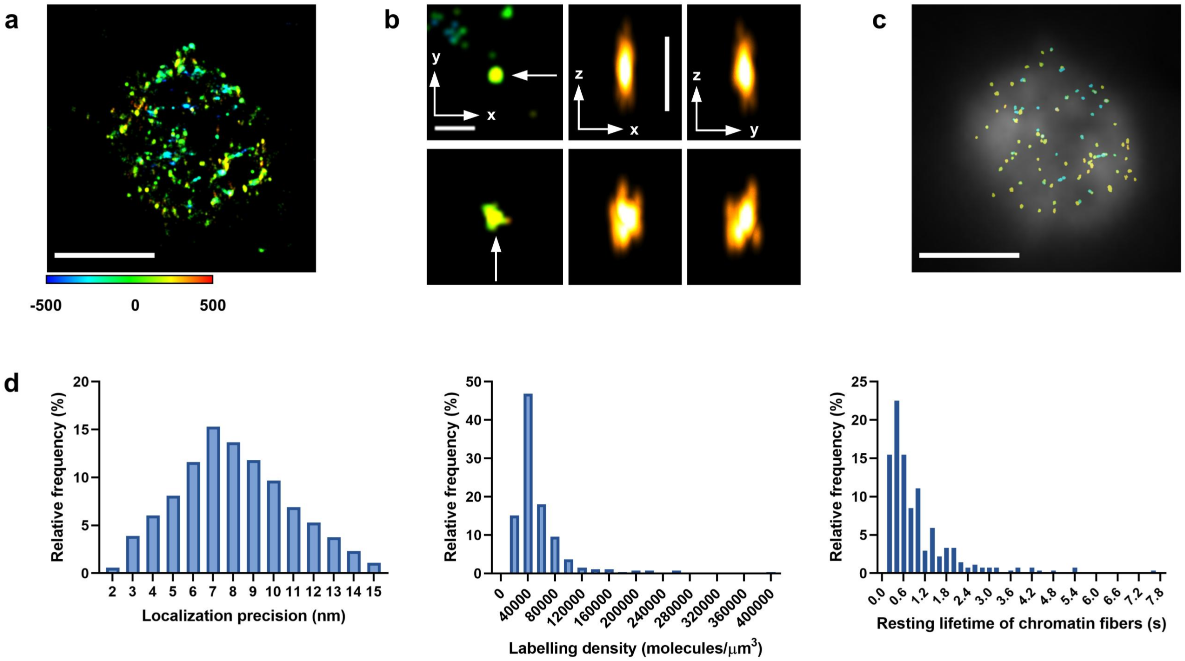


**Supplementary Fig. 12 Chromatin fibers in living chicken erythrocytes. a**, The image of a nucleus. Reconstructed with 5,000 frames (17.7 ms/frame). **b**, Chromatin fibers. FWHM (x, y, z) in nm (upper, lower): 25, 33, 84; 51, 47, 94. Localization precision (nm): 7.4, 8.5. **c**, The distribution of chromatin fibers (data from 5,000 frames). **d**, Localization precision, labeling density, and resting lifetime for identified fibers in 5,000 frames. Incubation with **6-HoeHESiR** (0.5 μM) for 10 min. Scale bar: 2 μm in **a** and **c**; 200 nm in **b**. Laser intensity: 0.95 kW/cm$^2$ at 656 nm. Axial position (nm) represented by RGB color depth coding.

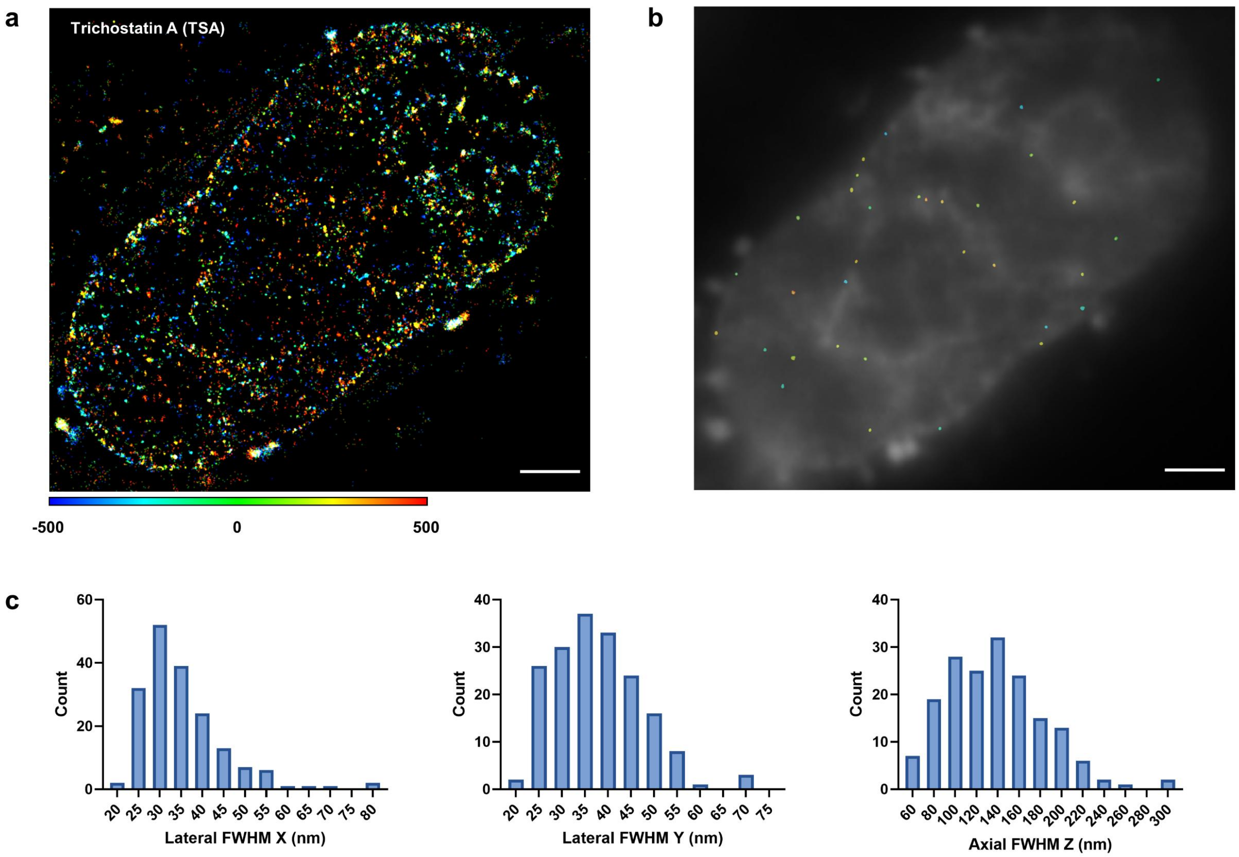


**Supplementary Fig. 13 Treatment with trichostatin A. a**, The nucleus of a treated HeLa cell. Reconstructed with 2,000 frames (17.7 ms/frame). **b**, The distribution of chromatin fibers (identified in 500 frames). **c**, 3D sizes of chromatin fibers from 5 cells. $N = 180$. Identified in 500 frames. Incubation with **6-HoeHESiR** (5 μM) for 30 min. Scale bar is 2 μm. Laser intensity: 3.2 kW/cm$^2$ at 656 nm. Axial position (nm) represented by RGB color depth coding.

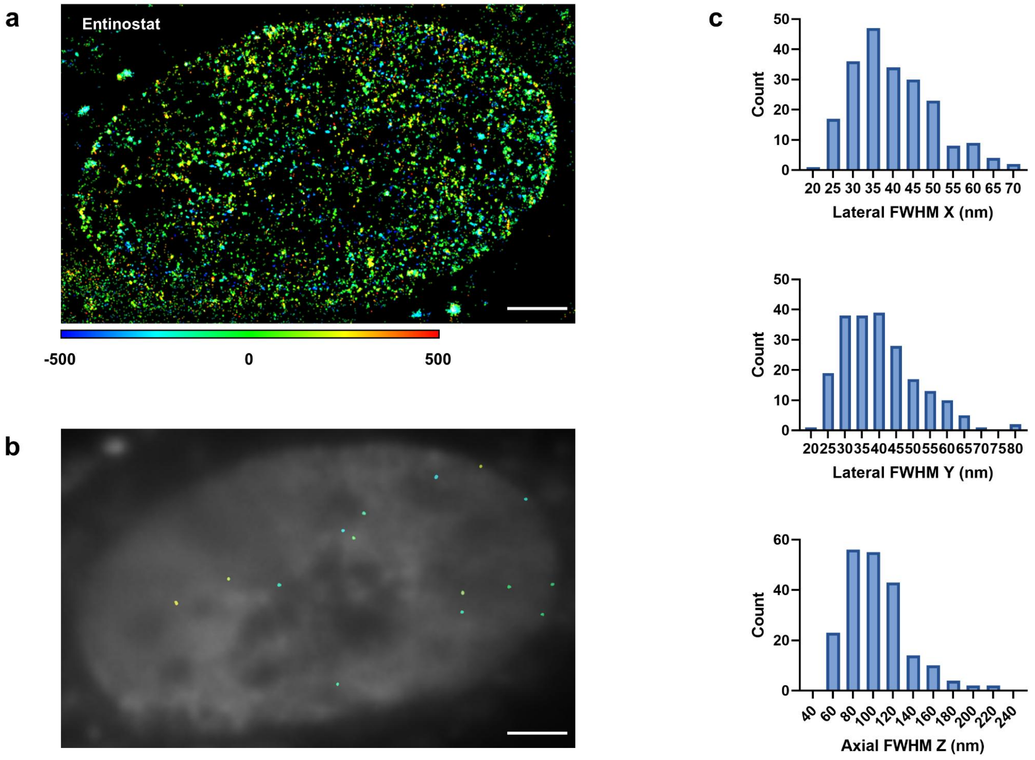


**Supplementary Fig. 14 Treatment with entinostat. a**, The nucleus of a treated HeLa cell. Reconstructed with 2,000 frames (17.7 ms/frame). **b**, The distribution of chromatin fibers (identified in 500 frames). **c**, 3D sizes of chromatin fibers from 5 cells. $N = 211$. Identified in 500 frames. Incubation with **6-HoeHESiR** (5 µM) for 30 min. Scale bar is 2 µm. Laser intensity: 3.2 kW/cm$^2$ at 656 nm. Axial position (nm) represented by RGB color depth coding.

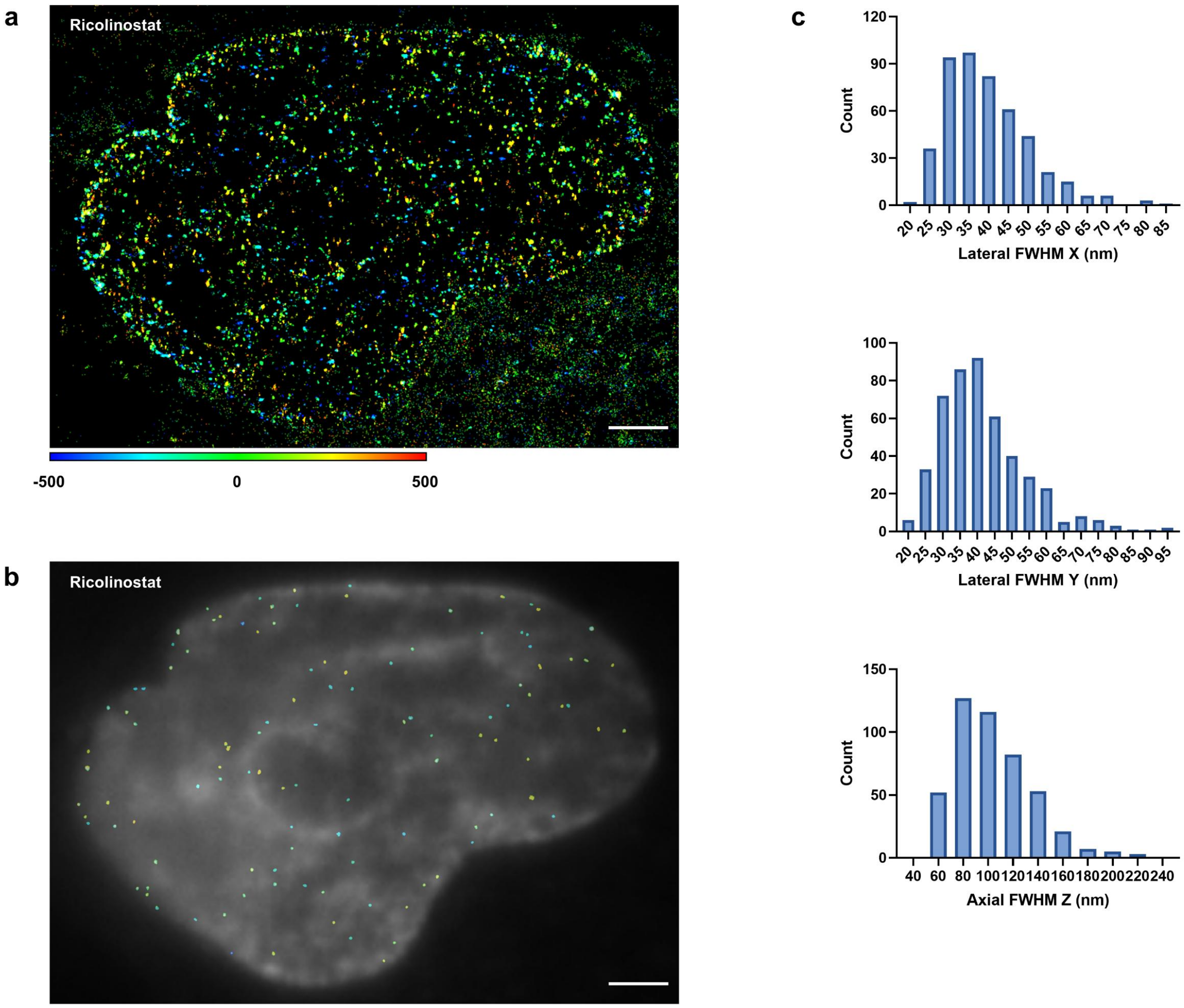


**Supplementary Fig. 15 Treatment with ricolinostat. a**, The nucleus of a treated HeLa cell. Reconstructed with 2,000 frames (17.7 ms/frame). **b**, The distribution of chromatin fibers (identified in 500 frames). **c**, 3D sizes of chromatin fibers from 5 cells. $N = 468$. Identified in 500 frames. Incubation with **6-HoeHESiR** (5 μM) for 30 min. Scale bar is 2 μm. Laser intensity: 3.2 kW/cm$^2$ at 656 nm. Axial position (nm) represented by RGB color depth coding.

#### Supplementary video legends

1. Self-blinking of **6-HoeHESiR** in a 3D SMLM experiment of *in vitro* reconstituted 30-nm chromatin fibers. Scale bar: 2 µm. 950 $W/cm^2$ laser intensity at 656 nm. Camera exposure at 17.7 ms/frame.
2. 3D rotation of an *in vitro* reconstituted chromatin fiber in Fig. 2g. Scale bar: 200 nm.
3. 3D rotation of another *in vitro* reconstituted chromatin fiber in Fig. 2g. Scale bar: 200 nm.
4. Self-blinking of **6-HoeHESiR** in a 3D SMLM experiment of a fixed HeLa cell. Scale bar: 5 µm. 3.2 $kW/cm^2$ laser intensity at 656 nm. Camera exposure at 17.7 ms/frame.
5. 3D rotation of a chromatin fiber in a fixed HeLa cell in Fig. 3c. Scale bar: 200 nm.
6. 3D rotation of another chromatin fiber in a fixed HeLa cell in Supplementary Fig. 7b. Scale bar: 200 nm.
7. Self-blinking of **6-HoeHESiR** in a 3D SMLM experiment of budding yeasts. Scale bar: 5 µm. 3.2 $kW/cm^2$ laser intensity at 656 nm. Camera exposure at 17.7 ms/frame.
8. 3D rotation of a chromatin fiber in a budding yeast in Fig. 3i. Scale bar: 200 nm.
9. 3D rotation of another chromatin fiber in a budding yeast in Fig. 3i. Scale bar: 200 nm.
10. Self-blinking of **6-HoeHESiR** in a 3D SMLM experiment of a live HeLa cell. Scale bar: 2 µm. 3.2 $kW/cm^2$ laser intensity at 656 nm. Camera exposure at 17.7 ms/frame.
11. 3D rotation of a chromatin fiber in a live HeLa cell in Fig. 4e. Scale bar: 200 nm.
12. 3D rotation of another chromatin fiber in a live HeLa cell in Fig. 4e. Scale bar: 200 nm.
13. Time-lapse video (z-projection) of dynamic chromatin structures in a living HeLa cell. 56 frames per image. 5 frames per sliding window. Scale bar: 2 µm. 3.2 $kW/cm^2$ laser intensity at 656 nm. Camera exposure at 17.7 ms/frame. Axial position is represented by RGB color depth coding.
14. Time-lapse video (yz-projection) of a chromatin fiber in Fig. 4e. 56 frames per image. 5 frames per sliding window. Scale bar: 200 nm. 3.2 $kW/cm^2$ laser intensity at 656 nm. Camera exposure at 17.7 ms/frame.

15. Time-lapse video (z-projection) of chromatin dynamics in a living HeLa cell with reduced phototoxicity. 56 frames per image. 5 frames per sliding window. Scale bar: 2 μm. 950 W/cm$^2$ laser intensity at 656 nm. Camera exposure at 17.7 ms/frame. Axial position is represented by RGB color depth coding.
16. Self-blinking of **6-HoeHESiR** in a 3D SMLM experiment of a living chicken erythrocyte. Scale bar: 2 μm. 950 W/cm$^2$ laser intensity at 656 nm. Camera exposure at 17.7 ms/frame.
17. 3D rotation of a chromatin fiber in a chicken erythrocyte in Fig. 5e. Scale bar: 200 nm.
18. 3D rotation of another chromatin fiber of a chicken erythrocyte in Fig. 5e. Scale bar: 200 nm.
19. Time-lapse video (z-projection) of chromatin dynamics in a living chicken erythrocyte with reduced phototoxicity. 56 frames per image. 5 frames per sliding window. Scale bar: 2 μm. 950 W/cm$^2$ laser intensity at 656 nm. Camera exposure at 17.7 ms/frame. Axial position is represented by RGB color depth coding.

### Chemical synthesis and characterization

#### General information

NMR spectra were acquired on Bruker 400 and 500 MHz spectrometers. High resolution mass spectra were acquired on Bruker maXis II and Thermo Scientific DFS. Preparative reverse phase HPLC was performed on a Waters preparative HPLC system with a C18 reverse phase column. All chemicals and reagents were purchased from commercial sources and used without further purification. Dess-Martin periodinane (DMP), dry *N*,*N*-diisopropylethylamine (DIEA), triethylamine and trifluoroacetic acid (TFA) were purchased from Energy Chemical. DIEA was purchased from TCI. $ClPh_3PCH_2OMe$ and Hoechst 33258 were purchased from J&K. NaHMDS was purchased from Alfa Aesar. $NaBH_4$ was purchased from Lancaster Synthesis. Chloromethyl methyl ether (MOMCl) was purchased from Sigma Aldrich. *s*-BuLi was purchased from Infinity Scientific. 4-Nitrophenyl chloroformate was purchased from Apollo Scientific. All solvents were either AR or HPLC grade. Dry dichloromethane ($CH_2Cl_2$) and dry tetrahydrofuran (THF) were obtained from a solvent purification system. Dry dimethylformamide (DMF) was purchased from Energy Chemical. Deuterated solvents $CDCl_3$ and $CD_3OD$ for NMR experiments were purchased from Cambridge Isotope Laboratories.

#### Synthetic route of 6-HESiR

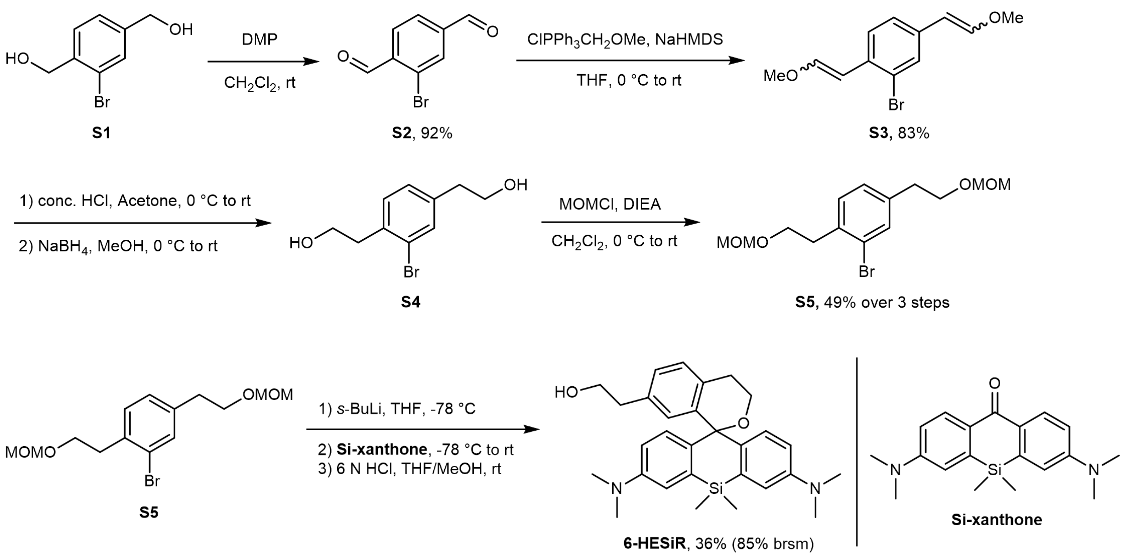

#### Procedures for making 6-HESiR

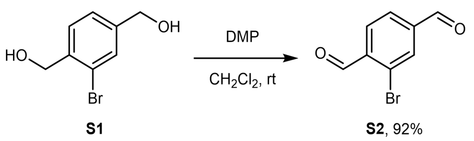

S1[1] (0.868 g, 4 mmol) and Dess-Martin periodinane (3.64 g, 8.4 mmol) were added to a 100 mL round-bottom flask, followed by $CH_2Cl_2$ (40 mL). The mixture was stirred at room temperature for 3 h. The milky suspension was quenched by saturated $NaHCO_3$ and extracted with $CH_2Cl_2$ three times. The combined organic phase was dried over $MgSO_4$, filtered and concentrated. The crude product was purified by flash column chromatography (10% to 20% ethyl acetate/hexane) to give the known compound **S2**[2] as a white solid (0.788 g, 3.7 mmol, 92% yield).

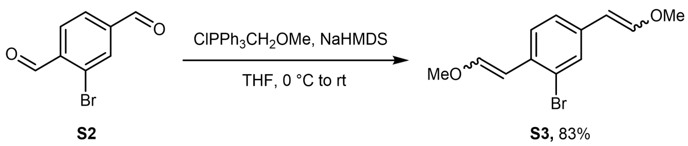

To a 100 mL round-bottom flask with $ClPh_3PCH_2OMe$ (1.58 g, 4.49 mmol, dried before reaction) under argon (Ar) was added THF (anhydrous, 7 mL). The flask was cooled in an ice/water bath with stirring under Ar. 2 M THF solution of NaHMDS (2.25 mL, 4.5 mmol) was added at 0 °C and the resulting mixture was stirred at the same temperature for 1 h. **S2** (0.318 g, 1.49 mmol) dissolved in THF (anhydrous, 4 mL) was transferred to the first flask. Bath was removed and the reaction mixture was warmed to rt and stirred overnight. On the next day, the reaction was quenched with sat. $NH_4Cl$, diluted with ethyl acetate (EA) and water, and extracted with EA three times. The combined organic phase was dried over $MgSO_4$, filtered and concentrated. The crude product was purified by flash column chromatography (15% $CH_2Cl_2$/hexane, then 5% EA/hexane) to give **S3** as a light-yellow oil (0.332 g, 1.23 mmol, 83% yield). Due to instability of **S3**, only $^1$H NMR spectrum was acquired. Due to complexity of the spectrum caused by two pairs of E/Z isomers, only integral ratios are provided. $^1$H

NMR (500 MHz, $CDCl_3$) δ 7.97–7.93, 7.81–7.78, 7.41–7.36, 7.25–7.22, 7.11–7.06 (3H, aromatic protons), 7.04–6.95, 6.22–6.20, 6.14–6.12, 6.09–6.04, 5.72–5.69, 5.59–5.56, 5.13–5.11 (4H, vinylic protons), 3.79–3.77, 3.72, 3.68 (6H, protons of two methyl groups).

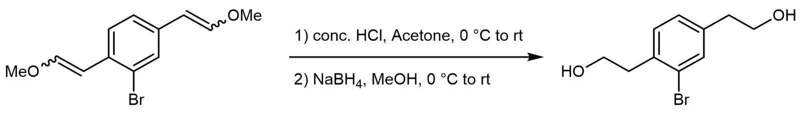


**S3** **S4**

**S3** (0.332 g, 1.23 mmol) was dissolved with acetone (13 mL) in a 100 mL round-bottom flask and the solution was cooled in an ice/water bath. Concentrated HCl (1 mL) was added to the solution and the mixture was stirred for 30 min at 0 °C. Then bath was removed, and the reaction was warmed to room temperature and stirred for another 1.5 h. The reaction was quenched with sat. $NaHCO_3$ and extracted three times with EA. The combined organic phase was washed with brine, dried over $MgSO_4$, filtered, and concentrated. Then methanol (14 mL) was added, and the round-bottom flask was cooled in an ice/water bath with stirring. $NaBH_4$ (0.375 g, 12.3 mmol) was added carefully (gas evolution) at 0 °C. The reaction mixture was stirred at the same temperature for 30 min. Then bath was removed, and the reaction was warmed to room temperature and stirred overnight (reaction time unoptimized). On the next day, saturated $NaHCO_3$ was added and EA was used to extract the mixture three times. The combined organic phase was washed with brine, dried over $MgSO_4$, filtered, and concentrated. The crude product was purified by flash column chromatography (2% to 4% $MeOH/CH_2Cl_2$) to give **S4** as a light-yellow viscous oil mixed with inseparable impurities, which was used for the next step without further purification. The crude product could also be directly used for the next step without flash column chromatography.

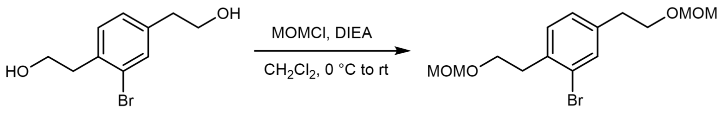


**S4** **S5,** 49% over 3 steps

To a 100 mL round-bottom flask with **S4** under Ar were added dry $CH_2Cl_2$ (12 mL) and dry DIEA (1.26 mL, 7.38 mmol) sequentially. The flask was cooled in an ice/water bath with stirring under Ar. MOMCl (0.52 mL, 6.15 mmol) was added dropwise at 0 °C. The reaction mixture was gradually warmed to room temperature and stirred overnight (reaction time unoptimized). On the next day, the reaction was quenched with water and extracted three times with $CH_2Cl_2$. The combined organic phase was dried over $MgSO_4$, filtered and concentrated. The crude product was purified by flash column chromatography (5% to 10% EA/hexane) to give **S5** as a colorless oil (200 mg, 0.6 mmol, 49% yield over 3 steps). $^1$H NMR (500 MHz, $CDCl_3$) δ 7.44 (d, $J$ = 1.4 Hz, 1H), 7.20 (d, $J$ = 7.8 Hz, 1H), 7.11 (dd, $J$ = 7.8, 1.4 Hz, 1H), 4.62 (s, 2H), 4.61 (s, 2H), 3.75 (t, $J$ = 7.5 Hz, 2H), 3.74 (t, $J$ = 7.3 Hz, 2H), 3.31 (s, 3H), 3.30 (s, 3H), 3.02 (t, $J$ = 7.1 Hz, 2H), 2.85 (t, $J$ = 6.8 Hz, 2H); $^{13}$C NMR (100 MHz, $CDCl_3$) δ 139.5, 136.1, 133.4, 131.2, 128.3, 124.8, 96.7, 96.6, 68.3, 67.0, 55.54, 55.48, 36.4, 35.7; HRMS (EI): calcd for $C_{14}H_{21}BrO_4$ ($M^+$): 332.0623, 334.0603, found: 332.0613, 334.0600.

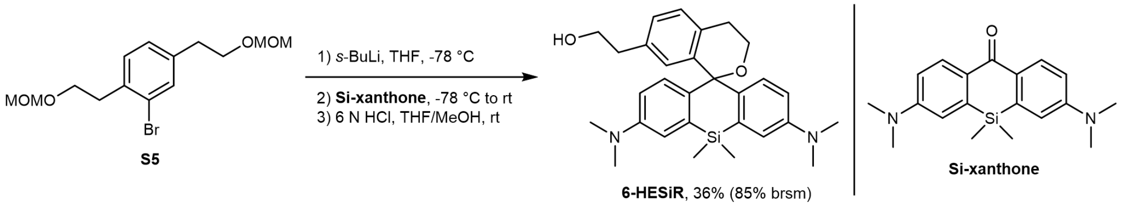


To a 100 mL round-bottom flask with **S5** (0.31 g, 0.93 mmol) under Ar was added THF (anhydrous, 4 mL). The solution was cooled in a dry ice/acetone bath with stirring. *s*-BuLi solution (0.97 mL, 0.96 M in hexane) was added dropwise at −78 °C and the resulting mixture was stirred for 3.5 h at the same temperature under Ar. Si-xanthone[3] (0.32 g, 0.99 mmol) was suspended in dry THF (sonication applied) and transferred to the first flask under Ar. This suspension/transfer operation was repeated several times (11 mL anhydrous THF in total) and THF (anhydrous, 2 mL) was added at the end to rinse the remaining residue. Upon completed transfer of Si-xanthone, the cold bath was removed, and the reaction was warmed to room temperature and stirred overnight (reaction time unoptimized). The flask was wrapped with aluminum foil to shield it from light. On the next day, the reaction was quenched with saturated $NH_4Cl$ and a small amount of 1 N HCl. The resulting deep blue mixture was diluted with water and extracted four times with $CH_2Cl_2$. The combined organic phase was dried over $Na_2SO_4$, filtered and concentrated. Then THF (4 mL) and MeOH (4 mL) were added, followed by 6 N HCl (10 mL). The reaction mixture was stirred at room temperature overnight (reaction time unoptimized). On the next day, the reaction mixture was diluted with water and quenched with solid $Na_2CO_3$ carefully (gas evolution). EA was added and the mixture was stirred vigorously until the deep blue color faded. Then the mixture was extracted with EA four times. The combined organic phase was washed with brine, dried over $MgSO_4$, filtered, and concentrated. The crude product was purified by flash column chromatography (20% EA/20% $CH_2Cl_2$/1% triethylamine/hexane to 50% EA/20% $CH_2Cl_2$/1% triethylamine/hexane) to give **6-HESiR** as a light blue foam (160 mg, 0.338 mmol, 36% yield). Based on recovered Si-xanthone (190 mg), the yield of **6-HESiR** based on recovered starting material (brsm) was 85%. $^1$H NMR (500 MHz, $CDCl_3$) δ 7.18 (d, $J$ = 7.8 Hz, 1H), 7.14 (d, $J$ = 7.8 Hz, 1H), 7.06 (s, 2H), 6.89 (s, 1H), 6.72 (d, $J$ = 8.8 Hz, 2H), 6.50 (d, $J$ = 8.7 Hz, 2H), 3.76 (t, $J$ = 6.6 Hz, 2H), 3.58 (t, $J$ = 5.2 Hz, 2H), 2.93 (s, 12H), 2.81 (t, $J$ = 5.2 Hz, 2H), 2.76 (t, $J$ = 6.5 Hz, 2H), 0.62 (s, 3H), 0.49 (s, 3H); $^{13}$C NMR (125 MHz, $CDCl_3$) δ 148.8, 139.3, 139.0, 138.3, 134.7, 134.6, 131.4, 130.5, 129.1, 127.3, 117.7, 112.1, 82.2, 63.9, 59.2, 40.6, 39.0, 29.2, 1.2, –2.7; HRMS (ESI): calcd for $C_{29}H_{36}N_2O_2Si$ ($[M + H]^+$): 473.2619, found: 473.2622.

**Synthetic route of 6-HoeHESiR**

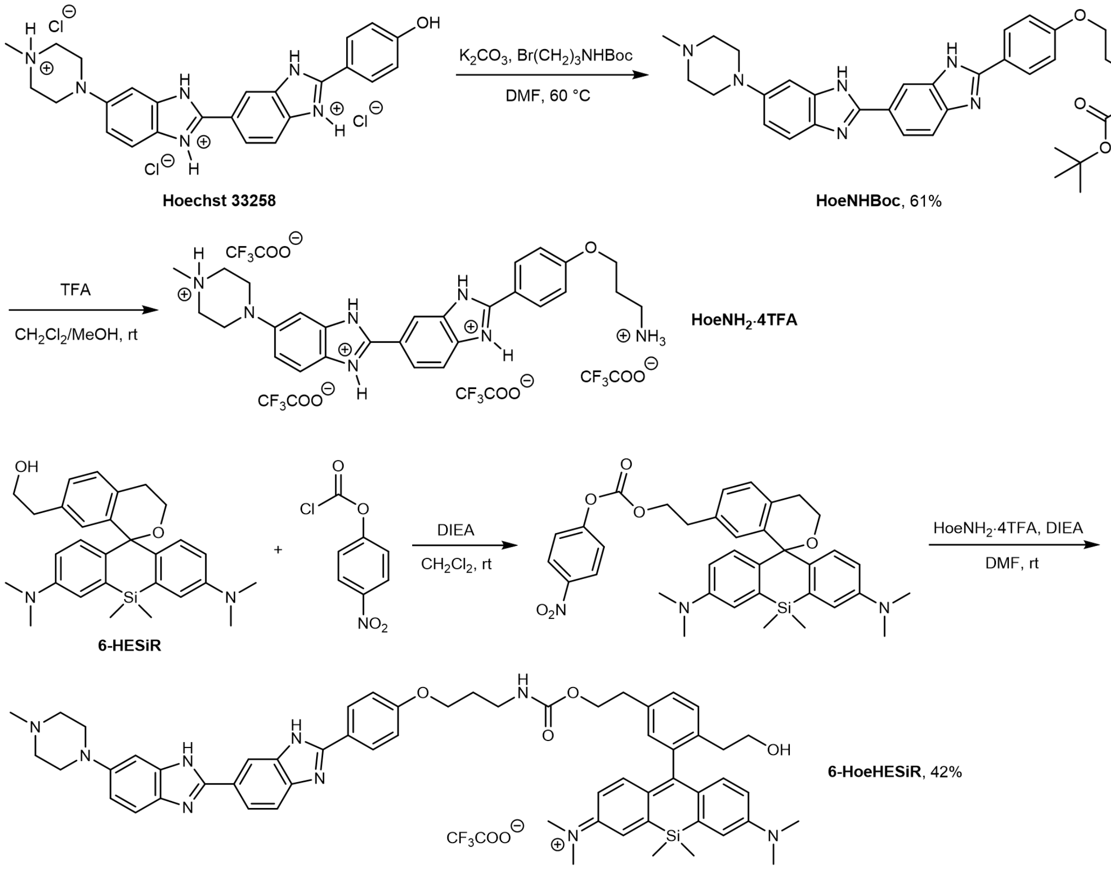


#### Procedures for preparation of 6-HoeHESiR

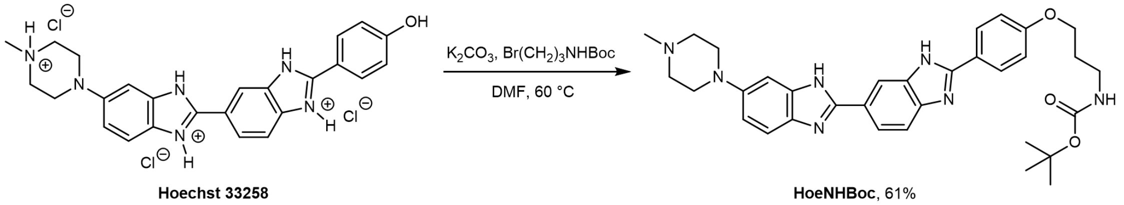


To a 25 mL round-bottom flask with **Hoechst 33258** (63 mg, 0.12 mmol), Br(CH$_2$)$_3$NHBoc[4] (56 mg, 0.236 mmol) and K$_2$CO$_3$ (49 mg, 0.354 mmol) was added DMF (3 mL). The suspension was stirred vigorously at 60 °C. The reaction mixture was shielded from light with aluminum foil. After two days (reaction time unoptimized), the reaction mixture was cooled to room temperature, diluted with methanol, and filtered. The filtrate was evaporated to dryness and the crude product was purified by flash column chromatography (0% to 10% MeOH/CH$_2$Cl$_2$) to give **HoeNHBoc** as a light yellow solid (42 mg, 0.072 mmol, 61% yield). $^1$H NMR (500 MHz, CD$_3$OD) δ 8.21 (s, 1H), 8.00 (d, *J* = 8.6 Hz, 2H),

7.91 (dd, $J$ = 8.5, 1.2 Hz, 1H), 7.65 (d, $J$ = 8.4 Hz, 1H), 7.50 (d, $J$ = 8.8 Hz, 1H), 7.13 (d, $J$ = 1.8 Hz, 1H), 7.03 (two doublets, $J$ = 8.6, 9.6 Hz, 2H + 1H = 3H), 4.03 (t, $J$ = 6.1 Hz, 2H), 3.29–3.26 (m, 4H), 3.24 (t, $J$ = 6.8 Hz, 2H), 2.93-2.87 (br, 4H), 2.55 (s, 3H), 1.94 (quintet, $J$ = 6.5 Hz, 2H), 1.43 (s, 9H); $^{13}$C NMR (125 MHz, $CD_3OD$) δ 162.5, 158.6, 155.3, 153.8, 149.1, 129.5, 125.5, 122.9, 122.5, 116.4, 116.0, 102.6, 80.0, 66.8, 55.8, 51.1, 45.3, 38.4, 30.7, 28.8; HRMS (ESI): calcd for $C_{33}H_{40}N_7O_3$ ($[M + H]^+$): 582.3187, found: 582.3201.

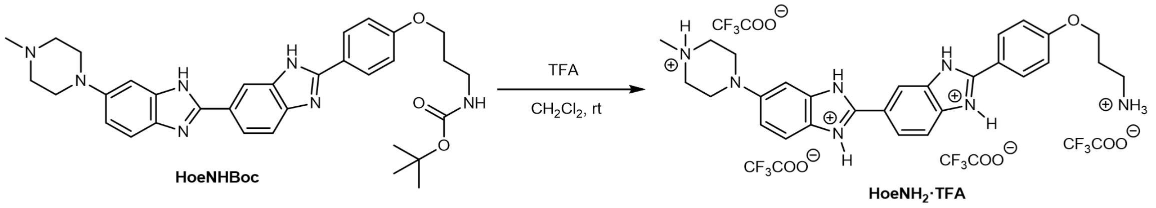


To a 10 mL round-bottom flask with **HoeNHBoc** (7.9 mg, 0.014 mmol) was added $CH_2Cl_2$ (0.5 mL) and TFA (0.5 mL). The suspension was stirred at room temperature for 1 h 10 min (reaction time unoptimized). Then toluene was added to the flask and the mixture was evaporated to dryness. **HoeNH₂·4TFA** salt was obtained as a brown oil and was used for the next step without further purification.

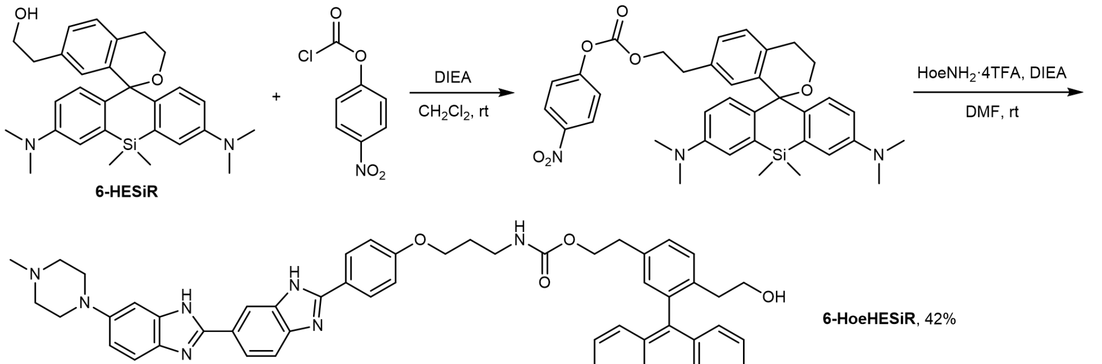


To a 10 mL round-bottom flask with **6-HESiR** (4 mg, 0.009 mmol) was added $CH_2Cl_2$ (0.4 mL) and DIEA (0.1 mL). 4-Nitrophenyl chloroformate (8.5 mg, 0.042 mmol) was added in one portion. The flask was sealed with a septum and stirred at room temperature overnight (reaction time unoptimized). On the next day, the reaction mixture was diluted with EA and quenched with water. The mixture was stirred vigorously for 15 min and then washed with water and brine. The organic phase was dried over $MgSO_4$, filtered, and concentrated. This mixed carbonate crude product was transferred to a 10 mL round-bottom flask, which was kept under Ar. DMF (anhydrous, 0.5 mL) and DIEA (anhydrous, 0.2 mL) were added to this flask. The resulting solution was added to the 10 mL round-bottom flask with **HoeNH$_2$·4TFA** salt under Ar. The first flask with mixed carbonate was rinsed with DMF (anhydrous, 0.3 mL). The reaction mixture was stirred at room temperature overnight under Ar. On the next day, the reaction mixture was passed through a 0.2 µm syringe filter and diluted with water and acetonitrile ($CH_3CN$). The crude product was purified by preparative reverse phase HPLC (30% to 70% $CH_3CN/H_2O$ with constant 0.1% TFA) to give **6-HoeHESiR** as a deep blue solid (4 mg, 0.004 mmol, 42% yield). $^1H$ NMR (500 MHz, $CD_3OD$) δ 8.37 (s, 1H), 8.07 (d, *J* = 8.5 Hz, 2H), 8.00 (d, *J* = 8.4 Hz, 1H), 7.85 (d, *J* = 8.5 Hz, 1H), 7.69 (d, *J* = 8.9 Hz, 1H), 7.42 (AB system, *J* = 8.9, 10.3 Hz, 2H), 7.37−7.27 (m, 4H), 7.12 (d, *J* = 9.8, 2H), 7.10 (d, *J* = 10.5 Hz, 2H), 7.03 (s, 1H), 6.76 (dd, *J* = 9.5, 1.7 Hz, 2H), 4.29 (t, *J* = 6.4 Hz, 2H), 4.06 (t, *J* = 5.8 Hz, 2H), 3.61-3.48 (br, 4H from the piperazine ring), 3.49 (t, *J* = 7.3 Hz, 2H), 3.42–3.29 (overlap with NMR solvent peak, 12H of the 4 methyl groups on 2 N atoms of the fluorophore structure), 3.28–3.24 (overlap with NMR solvent peak, 4H from the piperazine ring and 2H from a methylene group of the upper part of the fluorophore structure), 3.01 (s, 3H), 2.98 (t, *J* = 6.2 Hz, 2H), 2.52 (t, *J* = 7.3 Hz, 3H), 1.92 (quintet, *J* = 6.2 Hz, 2H), 0.60 (s, 3H), 0.57 (br, 3H); $^{13}C$ NMR (125 MHz, $CD_3OD$) δ 171.4, 164.0, 156.6, 151.0, 150.4, 143.7, 141.0, 138.8, 137.1, 132.1, 131.9, 131.6, 130.9, 129.9, 124.1, 122.9, 119.7, 117.2, 116.6, 115.9, 102.6, 67.8, 67.1, 63.8, 55.6, 44.4, 41.7, 39.6, 38.0, 36.9, 31.6, 31.5, 0.8, -0.7; HRMS (ESI): calcd for $C_{58}H_{66}N_9O_4Si$ ($M^+$): 980.5002, found: 980.5009.

**$^1H$ and $^{13}C$ NMR spectra of new compounds**

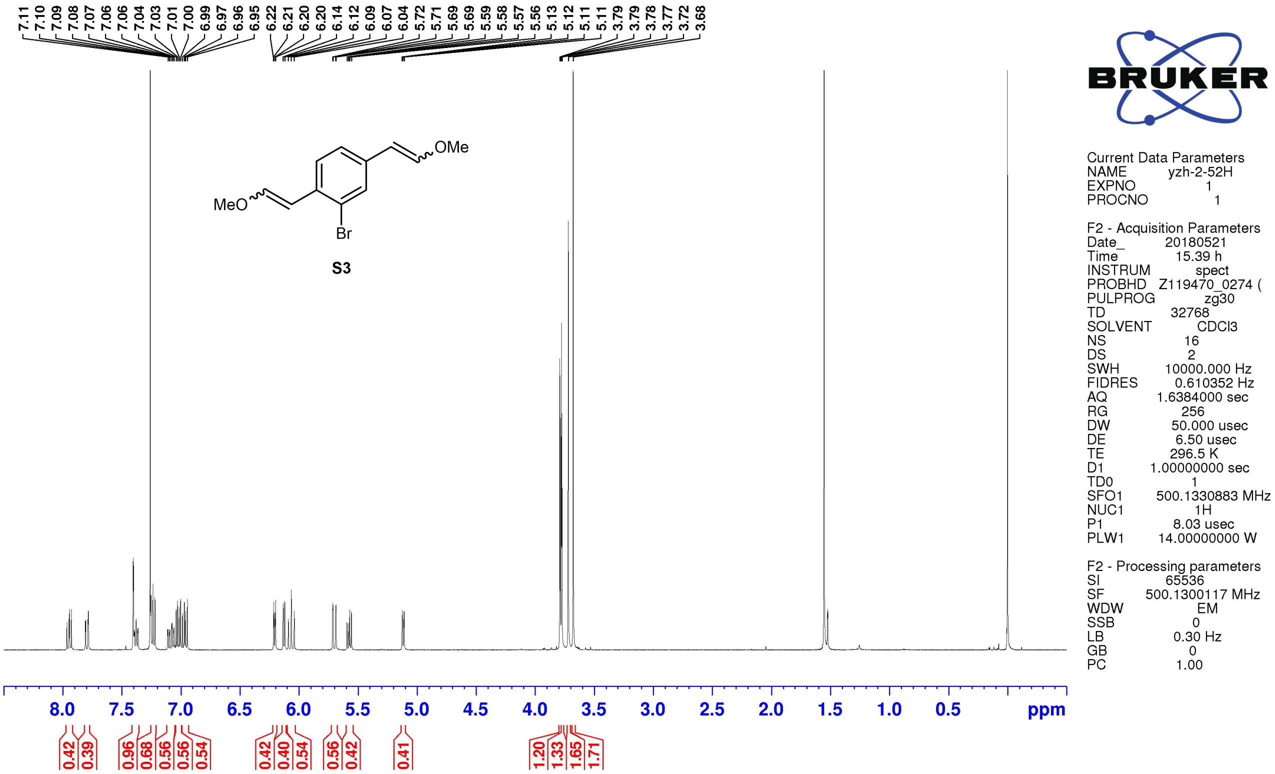
7.11 7.10 7.09 7.08 7.07 7.06 7.06 7.04 7.03 7.01 7.00 6.99 6.97 6.96 6.95 6.22 6.21 6.20 6.20 6.14 6.12 6.09 6.07 6.04 5.72 5.71 5.69 5.69 5.59 5.58 5.57 5.56 5.13 5.12 5.11 5.11 3.79 3.79 3.78 3.77 3.72 3.68
OMe
MeO
Br
S3
Current Data Parameters
NAME yzh-2-52H
EXPNO 1
PROCNO 1
F2 - Acquisition Parameters
Date_ 20180521
Time 15.39 h
INSTRUM spect
PROBHD Z119470_0274 (
PULPROG zg30
TD 32768
SOLVENT CDCl3
NS 16
DS 2
SWH 10000.000 Hz
FIDRES 0.610352 Hz
AQ 1.6384000 sec
RG 256
DW 50.000 usec
DE 6.50 usec
TE 296.5 K
D1 1.00000000 sec
TD0 1
SFO1 500.1330883 MHz
NUC1 1H
P1 8.03 usec
PLW1 14.00000000 W
F2 - Processing parameters
SI 65536
SF 500.1300117 MHz
WDW EM
SSB 0
LB 0.30 Hz
GB 0
PC 1.00
8.0 7.5 7.0 6.5 6.0 5.5 5.0 4.5 4.0 3.5 3.0 2.5 2.0 1.5 1.0 0.5 ppm
0.42 0.39 0.96 0.68 0.56 0.56 0.54 0.42 0.40 0.54 0.56 0.42 0.41 1.20 1.33 1.65 1.71

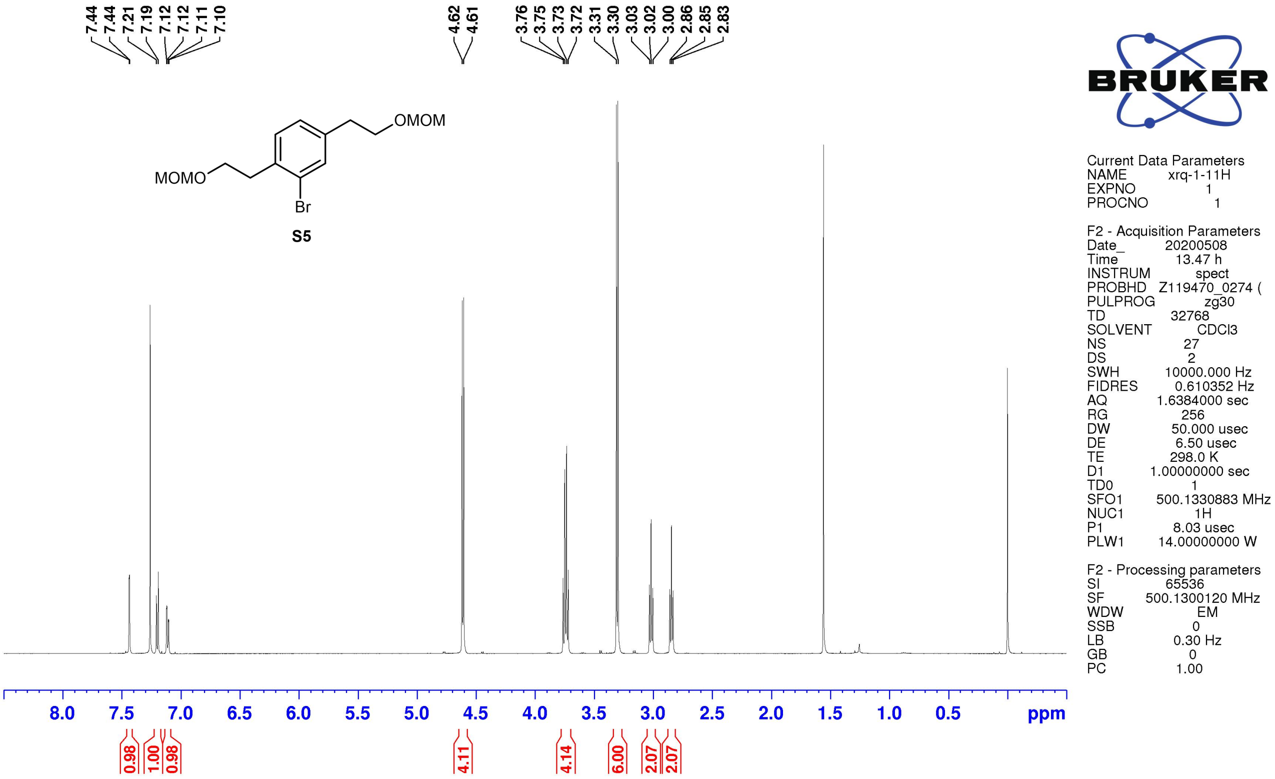
7.44 7.44 7.21 7.19 7.12 7.12 7.11 7.10 4.62 4.61 3.76 3.75 3.73 3.72 3.31 3.30 3.03 3.02 3.00 2.86 2.85 2.83
OMOM
MOMO
Br
S5
Current Data Parameters
NAME xrq-1-11H
EXPNO 1
PROCNO 1
F2 - Acquisition Parameters
Date_ 20200508
Time 13.47 h
INSTRUM spect
PROBHD Z119470_0274 (
PULPROG zg30
TD 32768
SOLVENT CDCl3
NS 27
DS 2
SWH 10000.000 Hz
FIDRES 0.610352 Hz
AQ 1.6384000 sec
RG 256
DW 50.000 usec
DE 6.50 usec
TE 298.0 K
D1 1.00000000 sec
TD0 1
SFO1 500.1330883 MHz
NUC1 1H
P1 8.03 usec
PLW1 14.00000000 W
F2 - Processing parameters
SI 65536
SF 500.1300120 MHz
WDW EM
SSB 0
LB 0.30 Hz
GB 0
PC 1.00
8.0 7.5 7.0 6.5 6.0 5.5 5.0 4.5 4.0 3.5 3.0 2.5 2.0 1.5 1.0 0.5 ppm
0.98 1.00 0.98 4.11 4.14 6.00 2.07 2.07

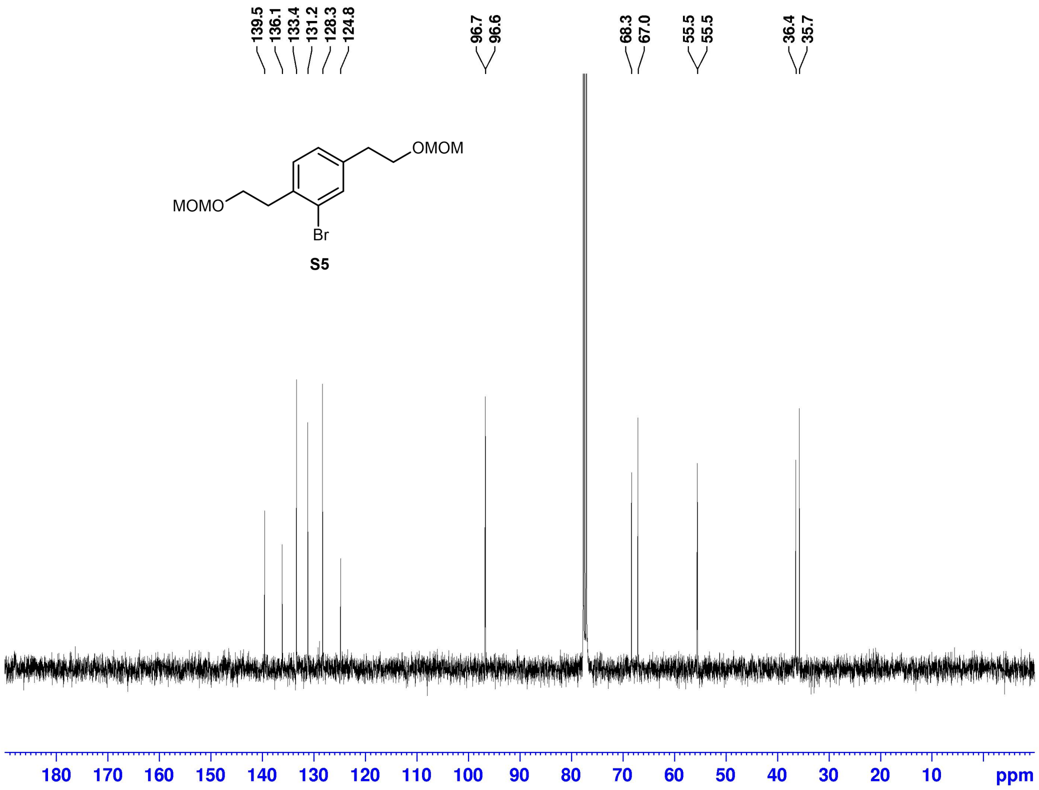


Current Data Parameters
NAME yzh-2-99C
EXPNO 1
PROCNO 1

F2 - Acquisition Parameters
Date_ 20180717
Time 10.27
INSTRUM spect
PROBHD 5 mm QNP 1H/1
PULPROG zgdc
TD 32768
SOLVENT CDCl3
NS 201
DS 0
SWH 25125.629 Hz
FIDRES 0.766773 Hz
AQ 0.6520832 sec
RG 26008
DW 19.900 usec
DE 20.00 usec
TE 0 K
D1 2.50000000 sec
d11 0.03000000 sec
MCREST 0 sec
MCWRK 0.01500000 sec

======== CHANNEL f1 ========
NUC1 13C
P1 9.50 usec
PL1 -5.00 dB
SFO1 100.6238364 MHz

======== CHANNEL f2 ========
CPDPRG[2 waltz16
NUC2 1H
PCPD2 80.00 usec
PL2 -4.00 dB
PL12 12.90 dB
SFO2 400.1320007 MHz

F2 - Processing parameters
SI 16384
SF 100.6127360 MHz
WDW EM
SSB 0
LB 1.00 Hz
GB 0
PC 1.40

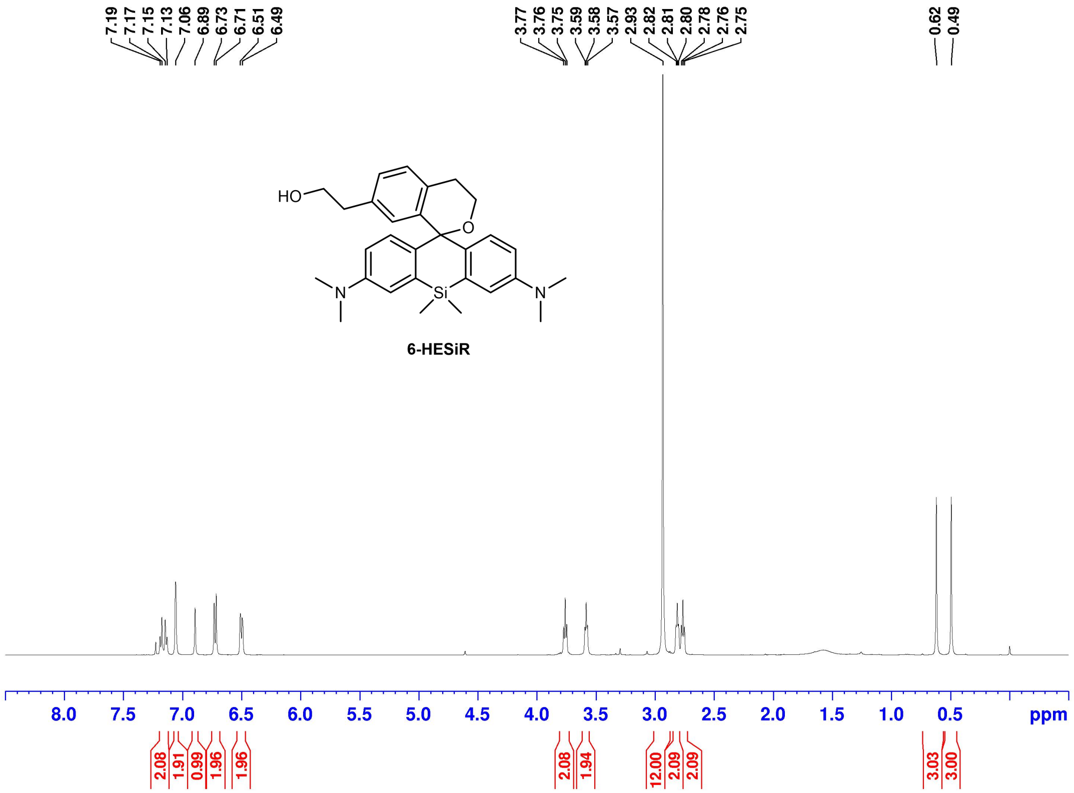


Current Data Parameters
NAME xrq-1-18H
EXPNO 5
PROCNO 1

F2 - Acquisition Parameters
Date_ 20200608
Time 13.43 h
INSTRUM spect
PROBHD Z119470_0274 (
PULPROG zg30
TD 32768
SOLVENT CDCl3
NS 32
DS 2
SWH 10000.000 Hz
FIDRES 0.610352 Hz
AQ 1.6384000 sec
RG 54.39
DW 50.000 usec
DE 6.50 usec
TE 295.4 K
D1 1.00000000 sec
TD0 1
SFO1 500.1330883 MHz
NUC1 1H
P1 8.03 usec
PLW1 14.00000000 W

F2 - Processing parameters
SI 65536
SF 500.1300291 MHz
WDW EM
SSB 0
LB 0.30 Hz
GB 0
PC 1.00

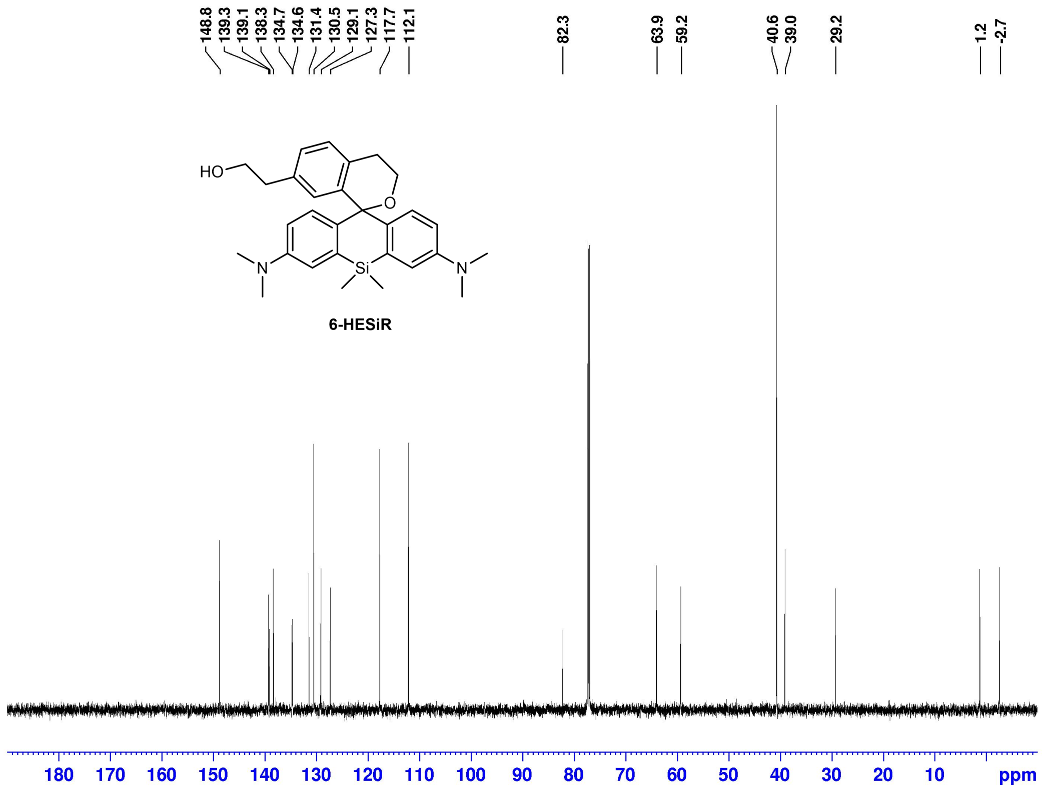


Current Data Parameters
NAME xrq-1-18C
EXPNO 1
PROCNO 1

F2 - Acquisition Parameters
Date_ 20200608
Time 14.26 h
INSTRUM spect
PROBHD Z119470_0274 (
PULPROG zgdc30
TD 32768
SOLVENT CDCl3
NS 30
DS 4
SWH 31250.000 Hz
FIDRES 1.907349 Hz
AQ 0.5242880 sec
RG 2050
DW 16.000 usec
DE 6.50 usec
TE 296.1 K
D1 2.00000000 sec
D11 0.03000000 sec
TD0 1
SFO1 125.7681547 MHz
NUC1 13C
P1 10.00 usec
PLW1 81.29100037 W
SFO2 500.1320005 MHz
NUC2 1H
CPDPRG[2 waltz16
PCPD2 80.00 usec
PLW2 23.04999924 W
PLW12 0.36015999 W

F2 - Processing parameters
SI 32768
SF 125.7577819 MHz
WDW EM
SSB 0
LB 1.00 Hz
GB 0
PC 1.40

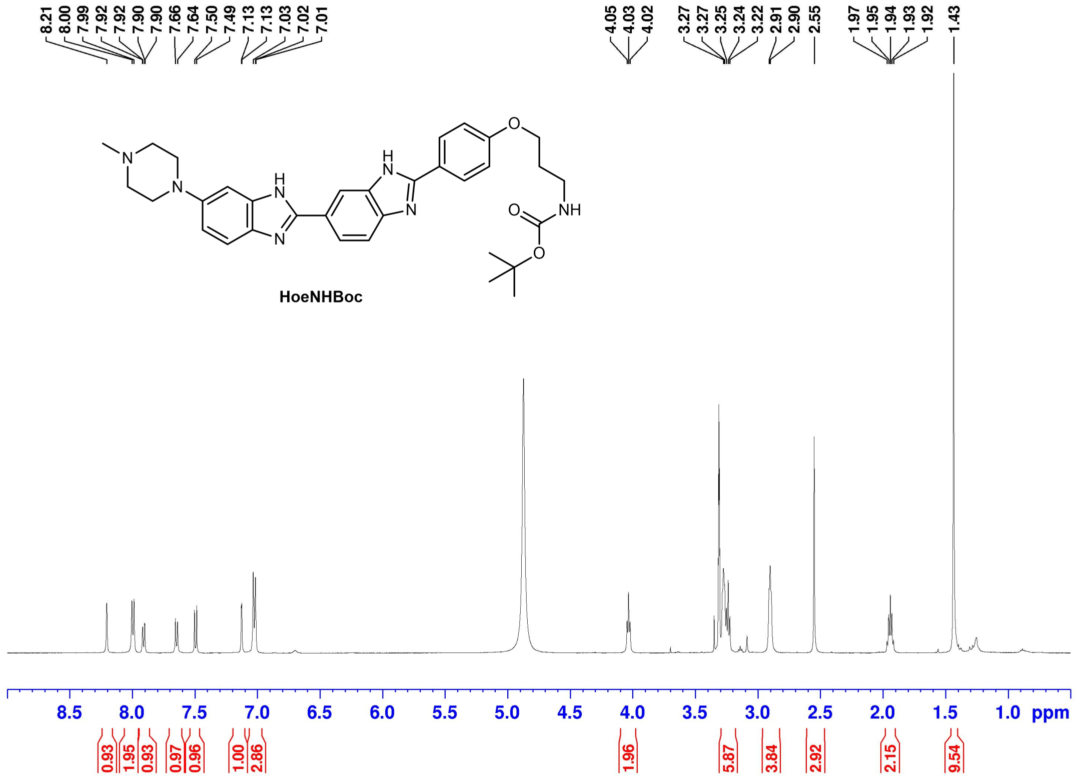


Current Data Parameters
NAME xrq-1-19H
EXPNO 2
PROCNO 1

F2 - Acquisition Parameters
Date_ 20200619
Time 14.11 h
INSTRUM spect
PROBHD Z119470_0274 (
PULPROG zg30
TD 32768
SOLVENT MeOD
NS 29
DS 2
SWH 10000.000 Hz
FIDRES 0.610352 Hz
AQ 1.6384000 sec
RG 117.5
DW 50.000 usec
DE 6.50 usec
TE 298.1 K
D1 1.00000000 sec
TD0 1
SFO1 500.1330883 MHz
NUC1 1H
P1 8.03 usec
PLW1 14.00000000 W

F2 - Processing parameters
SI 65536
SF 500.1300091 MHz
WDW EM
SSB 0
LB 0.30 Hz
GB 0
PC 1.00

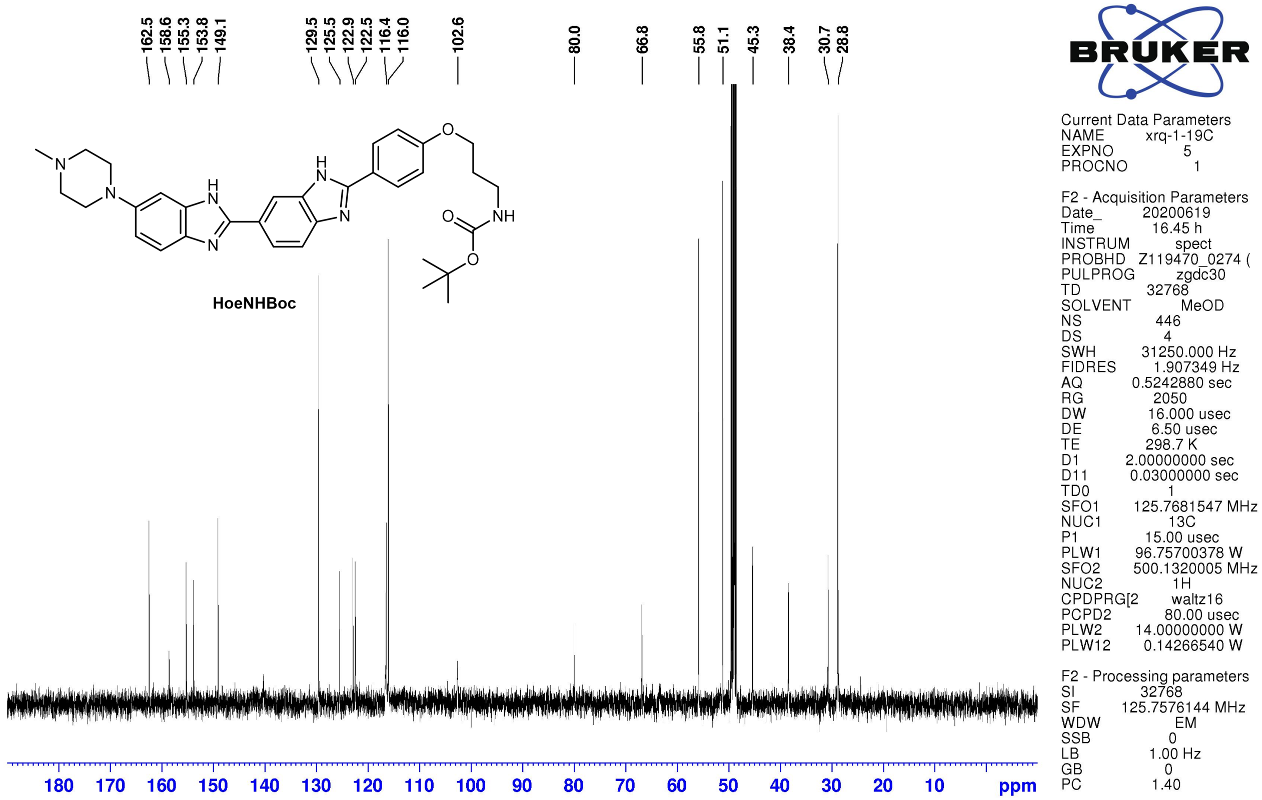
162.5
158.6
155.3
153.8
149.1
129.5
125.5
122.9
122.5
116.4
116.0
102.6
80.0
66.8
55.8
51.1
45.3
38.4
30.7
28.8
HoeNHBoc
BRUKER
Current Data Parameters
NAME xrq-1-19C
EXPNO 5
PROCNO 1
F2 - Acquisition Parameters
Date_ 20200619
Time 16.45 h
INSTRUM spect
PROBHD Z119470_0274 (
PULPROG zgdc30
TD 32768
SOLVENT MeOD
NS 446
DS 4
SWH 31250.000 Hz
FIDRES 1.907349 Hz
AQ 0.5242880 sec
RG 2050
DW 16.000 usec
DE 6.50 usec
TE 298.7 K
D1 2.00000000 sec
D11 0.03000000 sec
TD0 1
SFO1 125.7681547 MHz
NUC1 13C
P1 15.00 usec
PLW1 96.75700378 W
SFO2 500.1320005 MHz
NUC2 1H
CPDPRG[2 waltz16
PCPD2 80.00 usec
PLW2 14.00000000 W
PLW12 0.14266540 W
F2 - Processing parameters
SI 32768
SF 125.7576144 MHz
WDW EM
SSB 0
LB 1.00 Hz
GB 0
PC 1.40
180 170 160 150 140 130 120 110 100 90 80 70 60 50 40 30 20 10 ppm

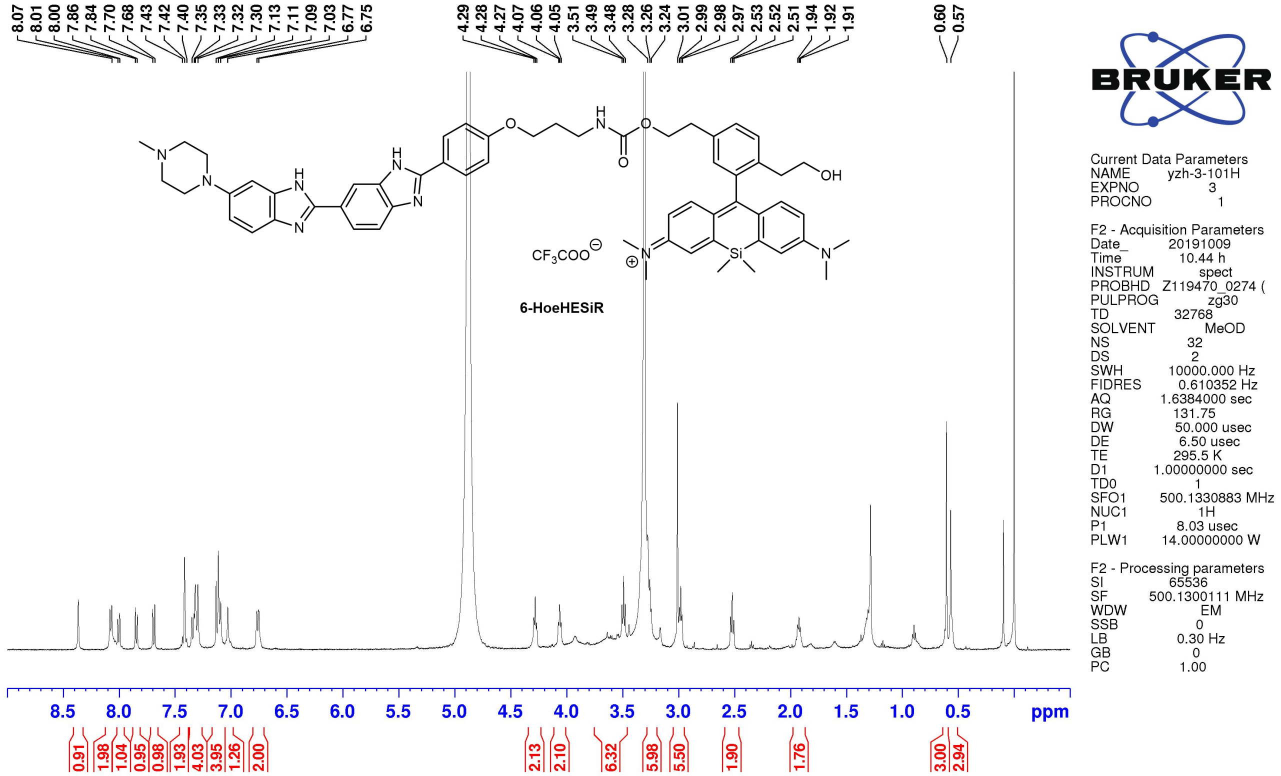
8.07
8.01
8.00
7.86
7.84
7.70
7.68
7.43
7.42
7.40
7.35
7.33
7.32
7.30
7.13
7.11
7.09
7.03
6.77
6.75
4.29
4.28
4.27
4.07
4.06
4.05
3.51
3.49
3.48
3.28
3.26
3.24
3.01
2.99
2.98
2.97
2.53
2.52
2.51
1.94
1.92
1.91
0.60
0.57
$CF_3COO^{\ominus}$
6-HoeHESiR
BRUKER
Current Data Parameters
NAME yzh-3-101H
EXPNO 3
PROCNO 1
F2 - Acquisition Parameters
Date_ 20191009
Time 10.44 h
INSTRUM spect
PROBHD Z119470_0274 (
PULPROG zg30
TD 32768
SOLVENT MeOD
NS 32
DS 2
SWH 10000.000 Hz
FIDRES 0.610352 Hz
AQ 1.6384000 sec
RG 131.75
DW 50.000 usec
DE 6.50 usec
TE 295.5 K
D1 1.00000000 sec
TD0 1
SFO1 500.1330883 MHz
NUC1 1H
P1 8.03 usec
PLW1 14.00000000 W
F2 - Processing parameters
SI 65536
SF 500.1300111 MHz
WDW EM
SSB 0
LB 0.30 Hz
GB 0
PC 1.00
8.5 8.0 7.5 7.0 6.5 6.0 5.5 5.0 4.5 4.0 3.5 3.0 2.5 2.0 1.5 1.0 0.5 ppm
0.91 1.98 1.04 0.95 0.98 1.93 4.03 3.95 1.26 2.00 2.13 2.10 6.32 5.98 5.50 1.90 1.76 3.00 2.94

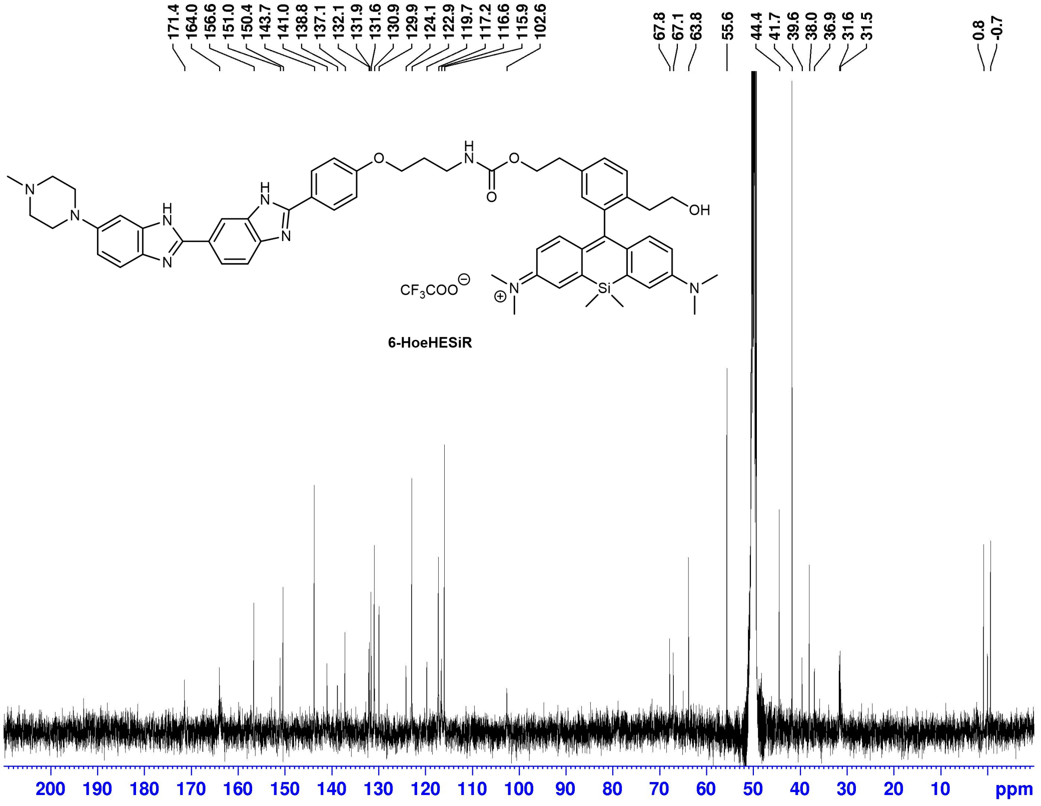


Current Data Parameters
NAME yzh-3-101
EXPNO 3
PROCNO 1

F2 - Acquisition Parameters
Date_ 20191009
Time 18.21
INSTRUM spect
PROBHD BBO 5mm Z3149/
PULPROG zgdc
TD 32768
SOLVENT MeOH
NS 15042
DS 1
SWH 31446.541 Hz
FIDRES 0.959672 Hz
AQ 0.5210112 sec
RG 9195.2
DW 15.900 usec
DE 6.00 usec
TE 0 K
D1 3.00000000 sec
d11 0.03000000 sec
MCREST 0 sec
MCWRK 0.01500000 sec

======== CHANNEL f1 ========
NUC1 13C
P1 11.00 usec
PL1 -2.00 dB
SFO1 125.7715724 MHz

======== CHANNEL f2 ========
CPDPRG[2 waltz16
NUC2 1H
PCPD2 100.00 usec
PL2 -6.00 dB
PL12 14.00 dB
SFO2 500.1312000 MHz

F2 - Processing parameters
SI 32768
SF 125.7575044 MHz
WDW EM
SSB 0
LB 1.20 Hz
GB 0
PC 1.00

### Materials and methods

#### Photophysical properties of 6-HESiR and 6-HoeHESiR

Absorbance was measured on a Shanghai INESA L6S UV-Vis spectrophotometer. Fluorescence emission was measured on an Edinburgh Instruments FS5 Spectrofluorometer. The hpDNA (5'-CGCGAATTCGCGTTTTCGCGAATTCGCG-3') used for fluorescence titration of **6-HoeHESiR** was purchased from Sangon Biotech. It was prepared by denaturation and annealing in the Tris-HCl buffer (50 mM Tris-HCl, 100 mM NaCl, pH = 7.4) with a digital dry bath. The dsDNA samples used for fluorescence titration studies were prepared from corresponding complementary oligonucleotides (Sangon Biotech) by denaturation and annealing in the same Tris-HCl buffer with a digital dry bath. The fluorescence titration of probe binding sites was performed with 0.5 μM **6-HoeHESiR** in the same Tris-HCl saline buffer by sequential addition of the stock solutions of hpDNA and dsDNA.

#### Configuration of the microscope for 3D SMLM experiments

All 3D SMLM experiments were performed on a Nano BioImaging SRiS 2.0 STORM Super-resolution Microscope[5] (https://cpos.hku.hk/imaging-and-flow-cytometry-core/imaging-equipment; company: INVIEW, http://www.inview-tech.com). 3D imaging was performed with astigmatism according to the reported approach[6]. The detailed configuration is listed below.

- Lasers: 647 channel: 656 nm (500 mW); 750 channel: 750 nm (300 mW)
- Microscope base: Nikon eclipse Ti-E
- XY stage: manual manipulator stage, piezo stage insert
- Objective lens: CFI Aopchromat TIRF 100XH, N.A. 1.49, W.D. 0.12 mm, cover glass correction: 0.13-0.21 mm
- 3D lens (for astigmatism): f = 1000.0 mm, Ø1", N-BK7 Mounted Plano-Convex Round Cylindrical Lens (LJ1516RM, Thorlabs)
- Detector: Andor iXon Ultra 897 EMCCD Camera; two-channel simultaneous acquisition with

320 × 160 pixels field of view

- Detection Band (647 channel, 656 nm laser): 672–712 nm (Model: FF01-692/40-25 made by Semrock)
- Dichoric mirror (for splitting between 647 and 750 channel): T760LPXR-UF2 (Chroma Technology Corp.)
- Software: Rohdea 2.0

**Nucleosome and chromatin reconstitution**

Recombinant histones H2A, H2B, H3, H4 and H1 were cloned and purified as previously described.[7] Reconstitution of histone octamers were performed as described previously.[8] Equimolar amounts of individual histones in unfolding buffer (7 M guanidinium HCl, 20 mM Tris-HCl, pH 7.5, 10 mM DTT) were dialyzed into refolding buffer (2 M NaCl, 10 mM Tris-HCl, pH 7.5, 1 mM EDTA, 5 mM 2-mercaptoethanol), and purified using Superdex S200 column. DNA templates of the 40–187bp DNA were cloned and purified as previously described.[8] Two single-stranded overhangs of DNA templates digested by BseYI enzyme were labeled with either dUTP-digoxigenin or dATP-biotin by Klenow reaction. Nucleosome samples were assembled using the salt-dialysis method as previously described.[8] Equimolar amounts of histone octamers and 40-187bp DNA templates were mixed in TEN buffers (10 mM Tris-HCl, pH 8.0, 1 mM EDTA, 2 M NaCl) and dialyzed for 17 hrs at 4 °C in TEN buffer, which was continuously diluted by slowly pumping in TE buffer (10 mM Tris-HCl, pH 8.0, 1mM EDTA) to lower concentration of NaCl from 2 M to 0.6 M. For histone H1 incorporation, histone H1 was added at this step and further dialyzed for 3 h. Samples were collected after final dialysis in HE buffer (10 mM HEPES, pH 8.0, 1 mM EDTA) for 4 h.

**Electron microscopy analysis**

Metal shadowing with tungsten for EM study was performed as described previously.[9] Reconstituted nucleosome samples (2 ng/μl) were fixed with 0.4% glutaraldehyde in HE buffer on ice for 30 min. After 2 mM spermidine was added, samples were applied to glow-discharged carbon-coated EM grids and incubated for 2 min and then blotted. Grids were washed stepwise in 0%, 25%, 50%, 75%, and 100% ethanol solution for 4 min at room temperature, air dried and shadowed with tungsten at an angle of 10° with rotation. For the negative staining, the chromatin samples in fixative solution were incubated on glow-discharged carbon-coated EM grids for 1 min. The excess sample solution was removed using filter papers. The grid was incubated in 2% Uranylacetate for staining for 30 s twice, blotted with filter papers and allowed to air-dry for several minutes. The prepared EM samples were examined using a FEI Tecnai G2 Spirit 120 kV transmission electron microscope.

**SMLM with spin-coated lambda DNA**

Lambda DNA (Thermofisher, SD0011) was dissolved in 1× TE50 buffer (pH = 7.5) to a concentration of 0.4 μg/mL and the solution was store at 4 °C. The stock solution of **6-HoeHESiR** was diluted in dd$H_2O$ to the concentration of 0.1 μM and the solution was stored at 4 °C. A confocal dish (cellvis, D35-10-1-N) was incubated with 0.1 mg/mL poly-L-lysine (Sigma) for 1 h. Then the well in the dish was washed with dd$H_2O$ three times and air-dried. The dish was fixed onto a spin-coater. The solution of lambda DNA (100 μL) was added to the dish. After 20 s of settling down, the spin speed was adjusted to 4000 rpm and kept for 60 s. During the spinning process, 2 mL dd$H_2O$ was added. After spin-coating, the solution of **6-HoeHESiR** (100 μL) was added to the dish for imaging. SMLM experiment was carried out on a home-built microscope with an Olympus IX73 base, a 100× NA = 1.5 oil-immersion objective and an Andor iXon897 EMCCD camera. 10,000 frames were acquired with laser intensity of 3.5 kW/$cm^2$ at 671 nm, 50 ms camera exposure, and EM gain of 30.

**3D SMLM of reconstituted nucleosomal arrays and 30-nm chromatin fibers**

A sample of *in vitro* reconstructed 30-nm chromatin fibers (40 × 187 bp with H1 and biotin label) and a sample of *in vitro* reconstructed nucleosomal arrays (40 × 187 bp without H1) were prepared. Streptavidin (1 mg, Sigma Aldrich, S4762-1MG) was dissolved in 1 mL PBS (1 mg/mL). 5 μL of this stock solution was diluted in 400 μL MQ water. In a 35 mm culture dish (Corning), this diluted solution was loaded onto an 18 mm round coverslip (Marienfeld precision cover glass thickness No. 1.5H) which has been sequentially coated with 3 μm microbeads (Sigma Aldrich, 79166-5ML-F) and poly-D-lysine (Sigma Aldrich, P0899-50MG, MW = 70,000−150,000). After 10 min, the solution was removed, and the coverslip was washed with MQ water. The sample of reconstructed 30-nm chromatin fibers (2.5 μL, 63.1 ng/μL) was added to 400 μL HE buffer (10 mM HEPES, 1 mM EDTA, pH = 7.5) containing 2.5 μM **6-HoeHESiR**. The resulting solution was loaded onto the coverslip with immobilized streptavidin. After 10 min, the solution was replaced with 380 μL HE buffer. 3D SMLM of reconstituted 30-nm chromatin fibers was carried out in HE buffer at room temperature by applying 950 $W/cm^2$ excitation laser intensity at 656 nm. Exposure time of EMCCD camera was 17.7 ms/frame. 5,000 frames were acquired for each field of view. The sample of reconstituted nucleosomal arrays (1 μL, 69 ng/μL) was added to 400 μL HE buffer containing 2.5 μM **6-HoeHESiR**. The resulting solution was loaded onto the coverslip coated with poly-D-lysine. After 10 min, the solution was replaced with 380 μL HE buffer. 3D SMLM of reconstituted nucleosomal arrays was carried out in HE buffer at room temperature using the same set of imaging parameters. Real-time drift correction was applied by the active locking function of Rohdea software using the microbeads coated on coverslip.

**3D SMLM of fixed HeLa cells**

HeLa cells were cultured in DMEM (Dulbecco's Modified Eagle Medium, Gibco) supplemented with phenol red, 10% heat-inactivated fetal bovine serum (Gibco) and 1% penicillin/streptomycin (Gibco), at 37 °C with 5% $CO_2$. A UV sterilized 18 mm round coverslip (Marienfeld precision cover glass thickness No. 1.5H) coated with 3 μm microbeads (Sigma Aldrich, 79166-5ML-F) was placed in a 35-mm culture dish (Corning) with coated face up. Cells were typically seeded at a density of $1.5 \times 10^5$

cells/mL the day before imaging. On the day of imaging, HeLa cells were fixed with 4% paraformaldehyde (PFA) in PBS for 15 min at rt. Then the cells were washed with PBS twice. The fixed cells were permeabilized with 1% Triton X-100 in PBS for 10 min at rt. Then the cells were washed with PBS twice. The fixed and permeabilized cells were stained with 8 μM **6-HoeHESiR** for 15 min in 1 mL PBS at rt. Then the cells were washed with PBS once. 3D SMLM of fixed HeLa cells was carried out in 380 μL PBS at room temperature by applying 3.2 kW/cm$^2$ excitation laser intensity at 656 nm. Exposure time of EMCCD camera was 17.7 ms/frame. 5,000 frames were acquired for each field of view. Real-time drift correction was applied by the active locking function of Rohdea software using the microbeads coated on coverslip.

**3D SMLM of fixed budding yeasts (*Saccharomyces cerevisiae* BY4741)**

The yeast strain *Saccharomyces cerevisiae* BY4741 used in this study is in S288C background. For routine cultures in this study, yeast cells were grown at 30 °C in YPD medium (1% yeast extract, 2% peptone, 2% dextrose). A single yeast colony from an agar plate was scraped for an overnight culture which was grown in 3 ml YPD at 280 RPM in a shaking incubator. The overnight yeast culture was diluted to $OD_{600} = 0.1$ in 3 ml YPD for mid-log phase culture. Yeast cells were collected at $OD_{600} = 0.4$–0.6 after at least two population doublings. The suspension (1 mL) of yeast cells was centrifuged and YPD medium was removed. The cells were washed with MQ water by resuspension and centrifugation twice. Then the cells were resuspended in 400 μL PBS and 200 μL of this suspension was loaded onto a microbead-coated coverslip having immobilized concanavalin A. After 15 min, the suspension was removed, and immobilized yeast cells were fixed with 4% PFA in PBS for 20 min at rt. Then the cells were washed with PBS once. The fixed cells were permeabilized with 1% Triton X-100 in PBS for 10 min at rt. Then the cells were washed with PBS twice. The fixed and permeabilized cells were stained with 8 μM **6-HoeHESiR** for 15 min in 1 mL PBS at rt. Then the cells were washed with PBS once. 3D SMLM of fixed HeLa cells was carried out in 380 μL PBS at room temperature by applying 3.2 kW/cm$^2$

excitation laser intensity at 656 nm. Exposure time of EMCCD camera was 17.7 ms/frame. 5,000 frames were acquired for each field of view. Real-time drift correction was applied by the active locking function of Rohdea software using the microbeads coated on coverslip.

**Light-sheet microscopy of living HeLa cells**

HeLa cells were cultured in DMEM (Dulbecco's Modified Eagle Medium, Gibco) supplemented with phenol red, 10% heat-inactivated fetal bovine serum (Gibco) and 1% penicillin/streptomycin (Gibco), at 37 °C with 5% $CO_2$. 5 mm coverslips (provided by Light Innovation Technology Ltd.) were sonicated in 75% ethanol for 10 min, air-dried and placed in a 35 mm culture dish. Cells were typically seeded at a density of 0.5 or 1 × $10^5$ cells/mL the day before imaging. On the day of imaging, Hela cells were incubated with **6-HoeHESiR** (5 µM for 30 min) and **6-HESiR** (2 µM for 30 min) in 1 mL DMEM at 37 °C with 5% $CO_2$. The cells were washed with DMEM once and placed back to the incubator with 1 mL DMEM in the dish. After 30 min, the cells were washed with DMEM once or twice before imaging. A coverslip with stained cells was placed in a sample holder and mounted. Live-cell 3D imaging was carried out in DMEM at 37 °C with 647-nm excitation laser and 30-ms camera exposure using a light-sheet microscope (Light Innovation Technology Ltd., LiTone LBS Light-sheet Microscope | litsite).

**Cytotoxicity assay**

HeLa cells were seeded in a 96-well plate at a density of 5000 cells/well and cultured in DMEM supplemented with phenol red, 10% heat-inactivated fetal bovine serum, 1% penicillin/streptomycin, and various concentrations of **6-HoeHESiR** at 37 °C with 5% $CO_2$. After 3 h, 10 µL of WST-1 reagent (Beyotime, C0036) was added to each well. After another 2 h, the absorbance readouts of each well were measured at 450 and 690 nm with a microplate reader (Thermo Varioskan LUX). Cell viability (%) data were obtained based on corrected absorbance values and a calibration curve of absorbance versus cell density.

**3D SMLM of living HeLa cells**

HeLa cells were cultured in DMEM supplemented with phenol red, 10% heat-inactivated fetal bovine serum and 1% penicillin/streptomycin, at 37 °C with 5% $CO_2$. A UV sterilized 18 mm round coverslip coated with 3 µm was placed in a 35-mm culture dish with coated face up. Cells were typically seeded at a density of 0.5 or 1 × $10^5$ cells/mL the day before imaging. On the day of imaging, Hela cells were incubated with 5 µM **6-HoeHESiR** for 30 min in 1 mL DMEM at 37 °C with 5% $CO_2$. The cells were washed with DMEM and placed back to the incubator with 1 mL DMEM in the dish. After 15–20 min, the cells were washed with DMEM before imaging. Live-cell 3D SMLM was carried out in 380 µL DMEM at room temperature by applying 3.2 kW/cm$^2$ or 950 W/cm$^2$ excitation laser intensity at 656 nm. Exposure time of EMCCD camera was 17.7 ms/frame. 5,000 frames were acquired for each field of view. Real-time drift correction was applied by the active locking function of Rohdea software using the microbeads coated on coverslip.

For treatments with HDAC inhibitors, HeLa cells were seeded on the day before imaging. Cells were treated with 200 nM TSA, 2 µM entinostat, and 1 µM ricolinostat for 20 h in DMEM, respectively. On the day of imaging, 3D SMLM experiments were carried out with the same experimental procedure and parameters as untreated cells.

**3D SMLM of living chicken erythrocytes**

20 µL chicken erythrocyte cell suspension (Innovative Research, 5% solution in Alsever's) was added to 1 mL DMEM containing 0.5 µM **6-HoeHESiR**. The suspended cells were incubated for 10 min in DMEM at 37 °C with 5% $CO_2$. The cells were centrifuged and DMEM was removed. Cells were washed with 1 mL HBSS (Hanks' Balanced Salt solution) and resuspended in 200 µL HBSS. The suspended cells in HBSS were loaded onto a microbead-coated coverslip having immobilized poly-D-lysine. The cells were allowed to settle down and attach to the coverslip for 5 − 10 min. Then the cell

suspension was replaced with 380 μL HBSS. Live-cell 3D SMLM was carried out in HBSS at room temperature by applying 950 W/cm$^2$ excitation laser intensity at 656 nm. Exposure time of EMCCD camera was 17.7 ms/frame. 10,000 frames were acquired for each field of view. Real-time drift correction was applied by the active locking function of the Rohdea software using the microbeads coated on coverslip.

**Chromatin image data analysis**

Super-resolved images were generated by FIJI[10] with ThunderSTORM[11] plugin using "Normalized Gaussian" reconstruction algorithm. The result table of acquired localizations for each field of view was generated by the Rohdea software and directly used for chromatin image data analysis. Magnification ratio is set as 10.6 to give a 10-nm pixel size in reconstructed images. Original pixel size is 106 nm and original field of view is 320 pixel (width) × 160 pixel (height). Axial position (nm) is represented by RGB color depth coding. Axial range is set as –500 to 500 nm. XZ and YZ 3D projection images of representative chromatin fibers were generated by the chromatin fiber analysis program and by Volume Viewer plugin in FIJI (orange hot pseudo-color). Blinking and time-lapse videos were produced by FIJI. 3D rotation videos of representative chromatin fibers were produced by Volume Viewer plugin in FIJI (orange hot pseudo-color).

We have developed a computer program for quantitative analysis of chromatin fibers. The program performs automated analysis of molecular clusters as potential chromatin fibers based on a set of user-defined parameters. The source code of the computer program is available at https://github.com/HKU-BAL/Chromatin-Fiber-Imaging. The flowchart of the analysis is presented in Supplementary Fig. 12. The program takes in an input table that contains "the coordinates of localized single molecules", "localization precision", "photon count", and "frame number". First, the input data is processed by a preprocessing module, which is applied for: 1) filtering out the imprecise localizations with the precision threshold set as 30 nm; 2) magnifying the X and Y coordinates with the ratio of 10.6; 3) grouping the single molecules based on Z-coordinates into 10 nm per slice; and 4) split the data table into 250 or 500 frames a group for ease of analysis. Second, the XY projection image of one frame interval data table is rendered by the "average shifted histogram" algorithm with lateral shift set as 2 and axis shift set as 4. The localized single molecules are assigned to respective clusters according to their 8-way pixel connectivity in the X and Y-axes. A noise removal module that applies the "gray value thresholding strategy" and "outlier Z slice removing by max z gap tolerance" is used. To preserve maximum information, for each instance, both the original cluster and the cluster after noise removal are kept for subsequent analyses. In the end, the XY, YZ, and XZ projection images of the identified clusters are rendered. For the identification of chromatin fibers with sufficient resolution and labeling density, we set up the following criteria: 1) continuous fibrous structure in YZ and XZ projection images; 2) labeling density ≥ 20,000 molecules/$\mu m^3$; 3) localization precision threshold of 15 nm; 4) maximum frame gap ≤ 40 frames; 5) maximum Z gap ≤ 50 nm. Long fiber diameter was the larger lateral FWHM value for each identified fiber. Short fiber diameter was the smaller lateral FWHM value for each identified fiber. Labeling density was calculated by the number of localizations for a chromatin fiber divided by its volume. The volume of a fiber is calculated by the total number of pixel cubes times the volume of each pixel cube, which is $10^{-6}$ $\mu m^3$. The resting/dwelling lifetime of a chromatin fiber was calculated by camera exposure 17.7 ms times the number of frames in which the localizations for that fiber were acquired.

The lateral and axial dimensions of a molecular cluster (a potential chromatin fiber) are estimated based on full width at half maximum (FWHM). To avoid potential bias in calculating FWHM, we adopted a "moving line" strategy. Using the estimation of the X dimension as an example, the process is illustrated in Supplementary Fig. 13a. The line moves on the cluster at the positions that cover the pixels at both the left and the right ends. The line width is set from 1 to the maximum that covers the whole cluster. Additionally, lines with too large FWHM fitting errors are discarded. The threshold is set as 6 for lateral FWHM and 30 for axial FWHM. The gray value submatrix within the line can be formulated as:

$$G_r = \begin{pmatrix} g_{i,j} & \cdots & g_{i,j+L} \\ \vdots & \ddots & \vdots \\ g_{i+w,j} & \cdots & g_{i+w,j+L} \end{pmatrix} \quad (1)$$

where $w$ denotes the line width. $L$ denotes the number of pixels horizontally of the cluster. $i, j$ represents the upper left position of the cluster. $g_{i,j}$ denotes the gray value at the position $(i,j)$

Then, the data points (**X**, **Y**) can be presented as:

$$\mathbf{X} = (x_0, x_1, \ldots, x_L) \quad (2)$$

$$x_k = (k - i) * a \quad (3)$$

$$\mathbf{Y} = (y_0, y_1, \ldots y_L) \quad (4)$$

$$y_k = \frac{\sum_{v=0}^{w} g_{i+v,k}}{w} \quad (5)$$

where $x_k$ denotes the distance between the $k^{th}$ position and the most left position of the line. $a$ represents the size of one pixel in the nanometer scale, which is 10 in this study. $y_k$ denotes the normalized gray value at $x_k$.

The FWHM of a molecular cluster is estimated by fitting a Gaussian curve:

$$y = y_0 + Ae^{-\frac{(x-x_c)^2}{2\omega^2}} \quad (6)$$

The parameters are initiated by:

$$y_0 = min\,(\mathbf{Y}) \quad (7)$$

$$A = max(\mathbf{Y}) - min\,(\mathbf{Y}) \quad (8)$$

$$x_c = \mathbf{X}_{argmax\,(\mathbf{Y})} \quad (9)$$

$$\omega = std(\mathbf{X}) \quad (10)$$

Then, the curve fitting is optimized by the Levenberg-Marquardt algorithm.

Finally:

$$FWHM = 2\sqrt{2ln2}\omega \quad (11)$$

$$FWHM_{err} = 2\sqrt{2ln2}\omega_{err}$$

The FWHM values are presented in two ways: 1) mean FWHM of all the lines, and 2) the FWHM with maximum line width.

To reduce potential “noise” signals close to the molecular clusters that might interfere with subsequent analysis, a “noise” removal module is utilized. The module consists of a “gray value thresholding” strategy and an “the outlier Z slice removing by max z gap tolerance” strategy (Supplementary information, Fig. 13b). The gray value thresholding is applied to XY, YZ and XZ projections of an identified molecule cluster. The key principle is “divide and conquer”. By removing the low gray value pixels, a large, connected cluster can be split into several smaller connected components and these components are analyzed by following the same process as the original cluster. On the other hand, the basic idea of the outlier Z slice removing strategy is to detect and discard the top and/or bottom localizations which are separated from nearest neighboring localizations by Z gaps.

**Quantification and statistical analysis**

Quantification and statistical analysis were performed with Microsoft Excel and Graphpad Prism 8. Statistical graphs were generated with Graphpad Prism 8.
